# Design of optical imaging probes by screening of diverse substrate libraries directly in disease tissue extracts

**DOI:** 10.1101/2020.06.08.140947

**Authors:** Martina Tholen, Joshua J. Yim, Katarzyna Groborz, Euna Yoo, Brock A. Martin, Nynke S. van den Berg, Marcin Drag, Matthew Bogyo

## Abstract

Fluorescently-quenched probes that are specifically activated in the cancer microenvironment have great potential application for diagnosis, early detection and surgical guidance. These probes are often designed to target specific enzymes associated with disease by direct optimization using single purified targets. However, this can result in painstaking chemistry efforts to produce a probe with suboptimal performance when applied *in vivo*. We describe here an alternate, unbiased activity-profiling approach in which whole tissue extracts are used to directly identify optimal peptide sequences for probe design. Screening of mouse mammary tumor extracts with a hybrid combinatorial substrate library (HyCoSuL) identified a combination of natural and non-natural amino acid residues that could be used to generate highly efficient tumor-specific fluorescently quenched substrate probes. The most effective probe is significantly brighter than any of our previously reported tumor imaging probes designed for specific proteases and robustly discriminates tumor tissue from adjacent healthy tissue in a mouse model of cancer. Importantly, although the probes were developed by screening mouse mammary tumor tissues, they are able to effectively distinguish human ductal carcinomas from normal breast tissue with similar reactivity profiles to those observed in mouse tissues. This new strategy simplifies and enhances the process of probe optimization by direct screening in a tissue of interest without any a priori knowledge of enzyme targets. It has the potential to be applied to advance the development of probes for diverse disease states for which clinical or animal model tissues are available.

## Introduction

There have been significant recent advances in the development of optical imaging probes for applications ranging from disease detection, diagnosis and monitoring to surgical guidance [1, 2]. Several fluorescent probes are currently in clinical trials for applications in image guided surgery (IGS), suggesting that it is likely that many types of surgery will use optical contrast in the near future. Generally, new classes of imaging probes are designed based on information about enzymes or receptors that are elevated or specifically expressed within a disease tissue of interest [3]. This information can then be used to tailor substrates or binding ligands to target a given disease state *in vivo*. While this approach has proven to be effective in some cases, it is often difficult to predict the *in vivo* performance of a given contrast agent that has been optimized for a single target *in vitro*. Probe design usually requires a labor-intensive, iterative process of optimization which may never result in molecules with a sufficient level of activity and/or selectivity for clinical applications. Therefore, approaches that would allow direct screening of libraries of potential contrast agents directly in the diseased tissue of interest would help to both accelerate the process of probe optimization while also allowing development of contrast agents against the most disease-relevant targets. Here we demonstrate that such an approach can be used with protease substrates to identify an optimal contrast agent for cancer imaging applications.

Due to the abundance of proteases in the human genome and their broad roles in the regulation of diverse disease pathologies, they make ideal target enzymes for development of imaging agents, diagnostics, and prodrugs [3, 4]. Typically, protease-activated agents are designed by identifying a peptide substrate sequence that is optimally cleaved to produce an optical signal or release an active drug. Elevation of protease activity in diseased tissue often results from inflammation [5] which is a hallmark not only of bacterial or viral infections but also of a multitude of chronic and age-related diseases, most notably cancer [6]. We and others have previously shown that protease activity that results from inflammation can be leveraged to visualize tumors [7, 8]. One prominent target for imaging probes of inflammation are lysosomal enzymes that are highly abundant in tumor-associated macrophages [9]. Among these enzymes, the cysteine cathepsins B, L, and S have been targets of a number of optical imaging probes [10–12]. Specifically, the fluorescently quenched substrate probe, 6QC, containing a dipeptide protease recognition sequence and a fluorophore/quencher pair, is effectively cleaved by cathepsins in tumor tissues and can be used for real-time imaging applications [13, 14]. This probe is a promising starting point for the development of intra-operative imaging contrast agents that can visualize tumor margins during surgery [14].

The process for developing proteolytically activated imaging agents or prodrugs usually starts with the selection of key proteases that are found in the tumor. After selecting the protease for targeting, its preferred substrate sequences are established, often using information from reported repertoires of endogenous substrates [15–17]. To further identify specificity determinants of a protease, it is often possible to screen diverse synthetic peptide substrate libraries using a purified, recombinant expressed version of the protease. This approach was originally developed for defining overall natural substrate specificity and predicting native protein substrates so it used only natural amino acid sequences [18]. Including diverse sets of non-natural amino acids into substrate libraries can enhance the chances of identifying highly efficient and specific protease substrates [19–22]. In fact, this approach has succeeded in identifying hybrid peptide substrates containing both natural and non-natural amino acids that show dramatically increased activity and selectivity for proteases within closely related families compared to substrates containing only natural amino acids [23, 24]. For example, highly specific covalent probes for related lysosomal cathepsins and apoptotic caspases were developed based on this approach [11, 25]. However, this general strategy still relies on predetermined information about the substrate specificity of the target protease, leading to a substantial bias of the libraries being screened. Furthermore, this approach essentially uses a single protease as a proxy for the overall proteolytic signature of a complex disease state for the purposes of probe design.

In this study, we show that it is possible to accelerate and enhance the process of probe design by direct screening of diverse substrate libraries in complex tumor tissues isolated from mouse mammary tissues. We used hybrid combinatorial substrate libraries (HyCoSuL) to identify multiple non-natural amino acid-containing peptide sequences that are optimally cleaved within these tissues. Using this information, combined with efforts to increase solubility and select for optimal cellular uptake, we identified a highly efficient fluorescently-quenched substrate probe that is suitable for direct *in vivo* imaging applications. This new probe is superior to our previously reported tumor imaging probes designed for lysosomal cathepsins. We also find that, although designed using mouse tumor tissues, the resulting imaging probe can identify human breast tumor tissues suggesting an overall conserved protease signature in mouse and human tumors. Overall, our results suggest that this approach for probe design has the potential to improve efforts to generate imaging probes for any disease application for which relevant whole tissue samples can be obtained.

## Results

### Proteolytic profiling of whole tumor tissue

Given our past success using fluorescent imaging probes for the detection and imaging of cancer, we chose to use tumor extracts from a mouse model of breast cancer for our proof of concept studies. Recent applications of highly diverse hybrid combinatorial substrate libraries (HyCoSuL) for protease substrate profiling have yielded selective inhibitors and activity-based probes (ABPs) for a number of classes of proteases including the lysosomal cysteine cathepsins [11, 23, 24]. We therefore chose to use HyCoSuLs for the tissue extract screening [26]. We performed proteolytic profiling of tissue lysates derived from tumor lesions induced in mice by mammary cell-driven expression of the Polyoma middle T oncogene [27, 28]. These mice carry a transgene that acts as an oncogene which is expressed under the control of the mouse mammary tumor virus long terminal repeat/promoter enabling mammary specific tumor formation. For the initial screening of tumor tissues, we decided to use material from a genetic tumor model rather than a cell injection model to ensure a better representation of intrinsic tumor development. Lysates from these tumors were prepared at both neutral and acidic pH to capture protease signatures of enzymes with diverse pH optima. We screened a HyCoSuL positional scanning library in which the P1 residue directly adjacent to the cleavage site is held constant as arginine to reduce complexity of the mixtures and because many proteases have P1 specificity for basic amino acids (**Fig. 1A, B** and **Fig. S1,2**). For each of the two sub-libraries, the ‘scanning’ P2 and P3 positions were varied through all the natural amino acids (minus cysteine and methionine) as well as a diverse set of 110 non-natural amino acids. The corresponding ‘non-scanning’ position contained an equimolar mixture of 18 natural amino acids plus norleucine (Mix). Comparing the cleavage activities of the substrate libraries at both neutral and acidic pH, we found that all substrates were uniformly more effectively cleaved at pH5.5 (**Fig. S3**). In general, we observed 10- to 20-fold higher cleavage activity at pH 5.5 so we used acidic pH for all further profiling studies. The pH 5.5 lysate profiling data for the libraries that scanned the natural amino acids revealed a preference for aliphatic and aromatic residues in the P2 position as well as a preference for leucine in the P3 position (**Fig. 1A, B**). This profile of natural amino acid specificity at acidic pH matched what would be expected for the lysosomal cathepsins, consistent with these enzymes being highly active in the tumor microenvironment. Further analysis of the diverse set of non-natural amino acids uncovered a number of residues with activity that was higher than the most optimal natural residues. At the P2 position the top residues contained mainly bulky aromatic sidechains (L-hSer(Bzl), Phe(3-I), Glu(O-Bzl), Ala(Bth)) as well as several straight chain aliphatic residues (2-Aoc, Abu, Nva). We therefore selected the top aromatic hit, hSer(Bzl), and the top aliphatic hit, (2-Aoc), for development of optical probes. For the P3 position, the aromatic Phenyl-glycine residue had the highest activity and was selected for use with the two optimal P2 residues.

**Fig. 1.**
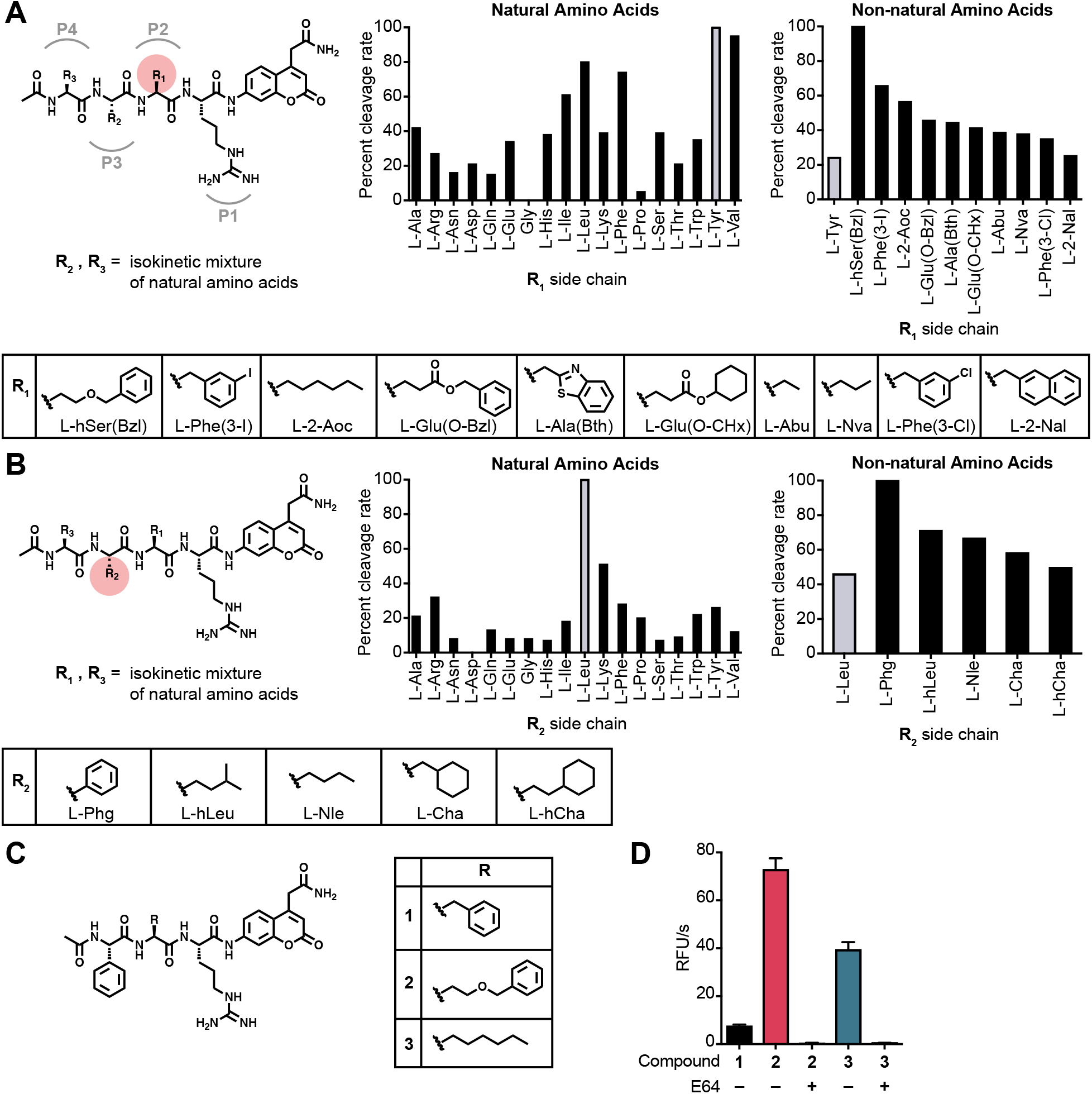
Proteolytic profiling of whole murine tumor tissue lysate with non-natural amino acid libraries. **(A)** Screening of the P2 scanning HyCoSuL library in mouse tumor tissue extract. The library contains arginine in P1, a variety of natural and non-natural amino acids in P2, and an isokinetic mixture of natural amino acids in P3 and P4. Plot of ACC cleavage relative to the most effectively cleaved residue is shown for natural amino acids (left) and the top non-natural amino acids with activity above the most optimal natural amino acid (right). Mean values of two tumor measurements were used. Structures of the non-natural amino acids from the plots are shown below. **(B)** Screening of the P3 scanning HyCoSuL library in mouse tumor tissue extract. The P3 scanning library was assayed exactly as in (A). **(C)** Structures of the ACC peptide substrates (**1**-**3**) synthesized based on optimal residues identified from the HyCoSuL screen in (A) and (B). **(D)** Plot of cleavage velocity of ACC substrates **1**-**3** in 4T1 murine breast cancer lysates with and without preincubation with cysteine cathepsin inhibitor E64 (10 μM). Mean value ± SD are plotted. n= 3.

In order to confirm that the selected amino acid residues could be used to make optimized imaging probes, we first synthesized the individual peptide substrates using the same ACC reporter that was used for the library screening (**Fig. 1C**). We synthesized peptide substrates that contained the P3 Phg and that had a P2 hSer(Bzl) (compound **2**) or 2-Aoc (compound **3**). We also included a substrate containing a P2 phenylalanine as a benchmark since it is the residue found on our previously reported cathepsin probe 6QC (compound **1**). This set of ACC substrates nicely matched the results from the positional scanning data with the P2 hSer(Bzl) substrate being most effectively cleaved and approximately 10-fold higher than the substrate containing the Phe residue. The 2-Aoc substrate showed about 50% of the activity of the hSer(Bzl) substrate **2** but remained 5-fold higher than the Phe substrate **1**. Importantly, preincubation with the covalent inhibitor E64 fully abrogated cleavage, suggesting the cysteine cathepsins as likely responsible for the majority of the proteolytic activity observed in the tumor extracts at acidic pH.

### Sequences selected from library screening can be converted to quenched substrate probes

To further validate the optimized substrate sequences identified from the library screening, we converted them to quenched substrate probes following the design of our previously described imaging probe, 6QC [13] (**Fig. 2A**). The 6QC probe (**4**) contains a Phe in P1 and has an N-terminal carboxybenzyl (Cbz) capping group that mimics the P3 residue. We therefore synthesized probes **5**-**8** containing a Phg in P3, either hSer(Bzl) or 2-Aoc in the P2 position and either a Cbz or an acetyl group as an N-terminal cap (**Fig. 2A**). All probes contained a Cy5 fluorophore attached to the peptide backbone via a six-carbon diamine linker that is paired with a QSY21 quencher attached to the P1 lysine sidechain amine. We tested each of the fluorescently quenched probes in murine tumor lysate at pH 5.5 and found that all of the substrate probes optimized from the library screening (**5**, **6, 7** and **8**) had increased activity compared to the original 6QC probe (**4**) (**Fig. 2B**). Interestingly, the acetylated versions of both probes (**6** and **8**) were more effectively processed than their counterparts containing the bulky Cbz capping group. Consistent with our library screening results, we found that probes with P2 hSer(Bzl) (**5** and **6**) were more effectively cleaved than both the original 6QC with a P2 Phe and the corresponding P2 2-Aoc containing probes. The most effective probe containing the acetate cap and the P2 hSer(Bzl) was approximately 65-fold more active in the tumor lysate than 6QC (**4**), and the corresponding carboxybenzyl capped probe, **5** was roughly 9 times more effective than 6QC **(4)**.

**Fig.2:**
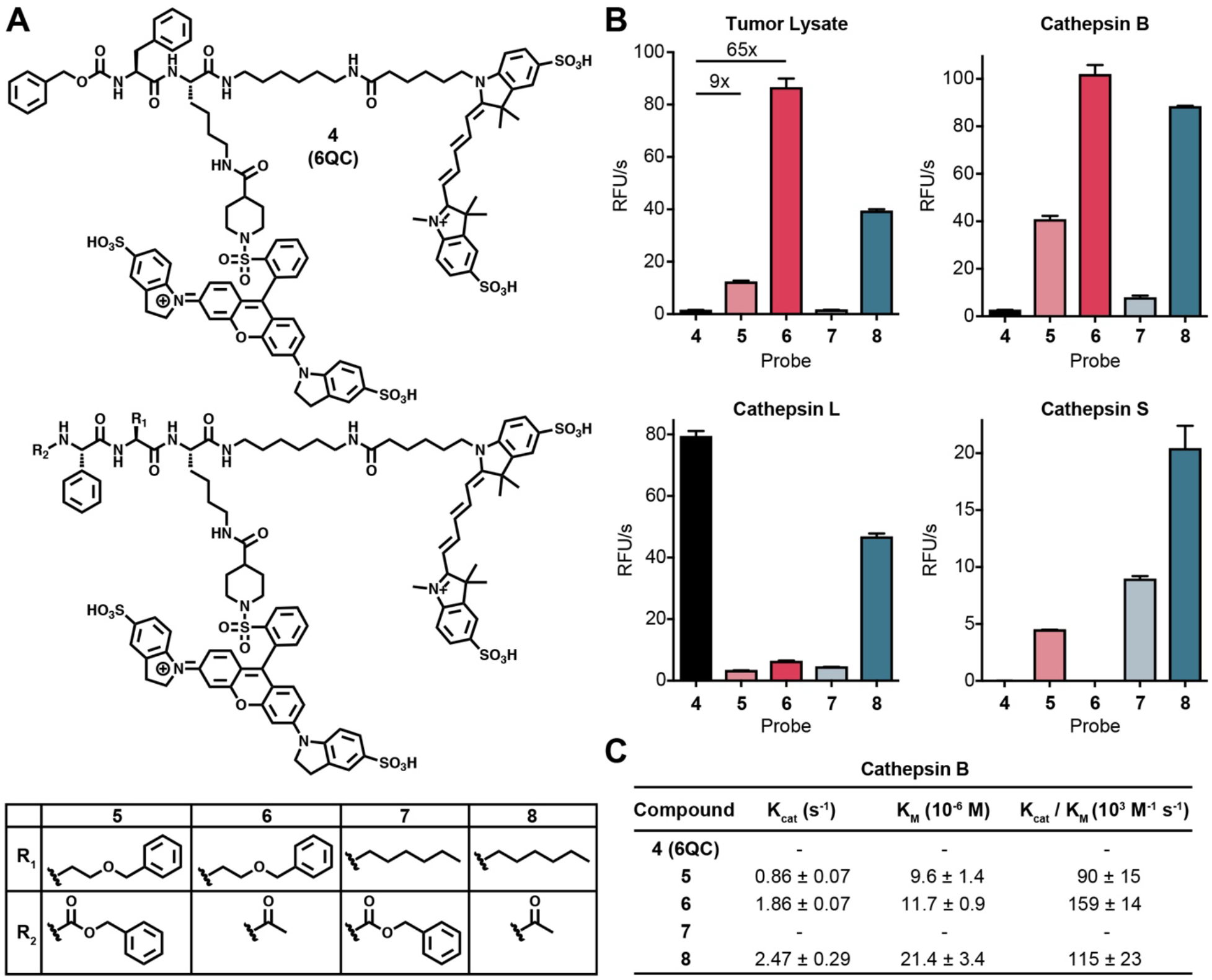
Conversion of substrate screening specificities into fluorescently quenched imaging probes. **(A)** Structure of the original fluorescently-quenched substrate probe 6QC compared to probes **5**-**8** that contain the optimized P2, P3 and N-terminal capping groups. **(B)** Cleavage rates of probes **4-8** when incubated with 4T1 murine breast tumor lysate at pH 5 (top left) or with purified cathepsin B (top right), cathepsin L (bottom left) or cathepsin S (bottom right). Values are plotted as mean value ± SD. n=3. **(C)** Kinetic parameters for the indicated probes for purified cathepsin B at pH 5. Values could not be obtained for probes **4** and **7** due to overall low activity.

Given our findings that the majority of enzyme activity responsible for processing the optimized probes in tissue extracts was likely due to cysteine cathepsins, we tested each of the probes for cleavage by recombinant versions of the three main cysteine cathepsins found in tumor associated macrophages (Cathepsins B, S and L). This analysis revealed that probes **5** and **6** were predominantly processed by cathepsin B while the related probes **7** and **8** were processed to varying degrees by all three cathepsins (Fig. 2B). In contrast, the original substrate probe 6QC (**4**) was only weakly processed by cathepsins B and S but was highly efficiently processed by cathepsin L. These results suggest that cathepsin B is the dominant proteolytic activity in the tumor tissue extracts at pH 5.5. We therefore measured the kinetic parameters of the three most active probes for processing by cathepsin B (**Fig. 2C**). The catalytic efficiency for **5**, **6**, and **8** is in the range of 90 to 160 10^3^M^−1^ s^−1^. This is similar to the reported efficiency of 6QC (**4**) for cleavage by cathepsin L which was reported to be 87 10^3^M^−1^ s^−1^ [13].

### Performance of probes in cells depends on permeability and solubility

Given the consistently higher activity of the probes containing the P2 hSer(Bzl), we focused on probes **5** and **6** for further testing and optimization. To investigate the ability of the optimized probes to be activated within cells, we measured the efficiency of cleavage of each probe by flow cytometry when added to immortalized macrophages (**Fig. 3A**). Quantification of the results revealed that the Cbz-capped probe **5** was more than two-fold more effectively cleaved than either the original 6QC (**4**) or the acetylated probe **6** (**Fig. 3B**). The overall low level of activity of probe **6** was surprising considering that it was the most effectively cleaved probe in the tumor tissue lysates as well as in the lysate of the immortalized macrophages (**Fig. S4**). We reasoned that this reduced activity for the acetyl capped probe could be the result of removal of the acyl group, resulting in a probe that was poorly cleaved by cathepsins. However, analysis of the activity of a free amino version of the probe excludes this explanation as it retained full activity for processing by cathepsins but also showed poor activity when applied to cells (**Fig. S5**).

**Fig. 3:**
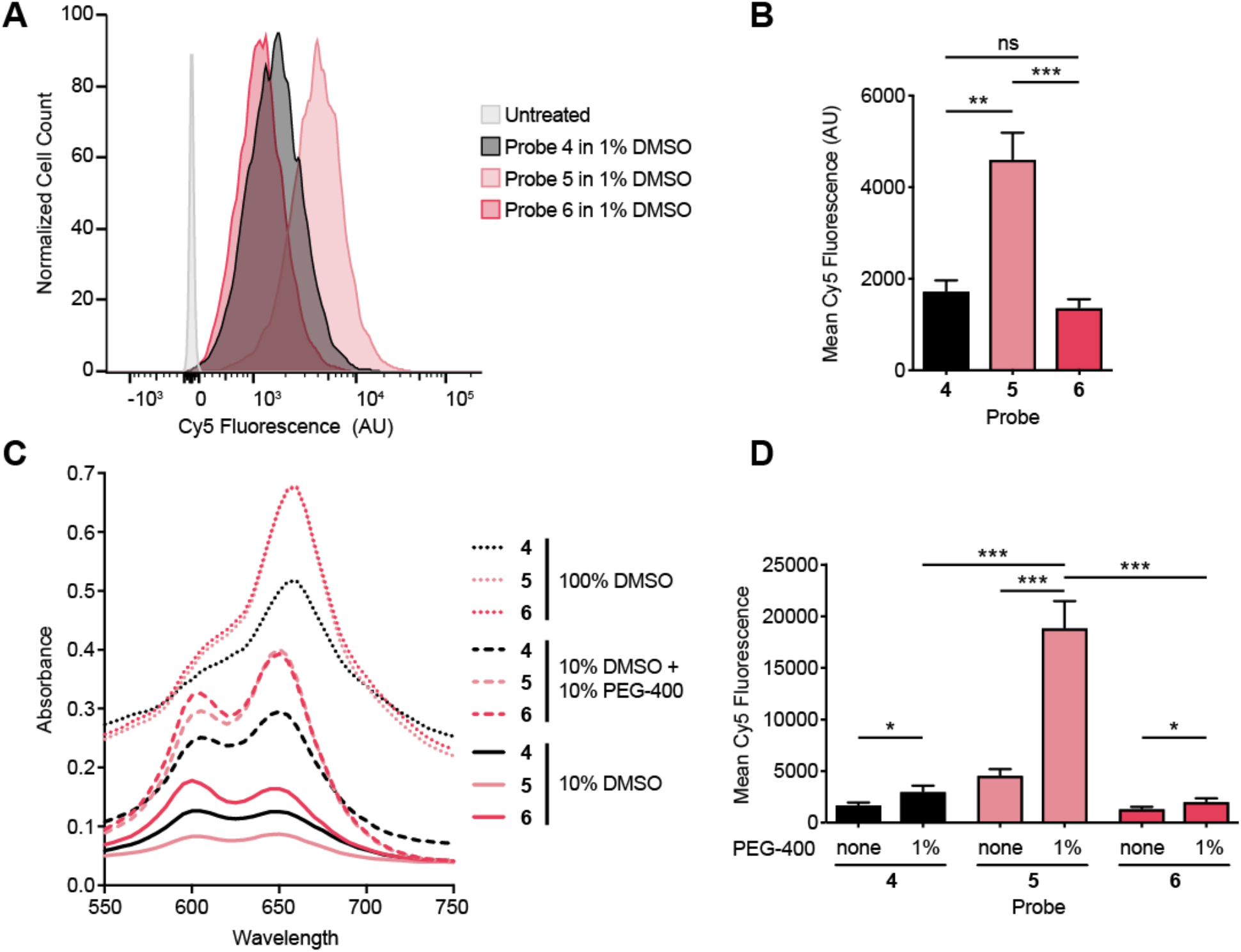
Further validation and optimization of quenched substrate probes using a cell-based assay. **(A)** Histogram of flow cytometry data for immortalized macrophages incubated with probes **4**-**6** (3 h incubation, 1 μM compound). **(B)** Quantification of fluorescent signals in (A) (3 h incubation, 1 μM compound). Mean value ± SD. n=3. Student’s t-test. **(C)** Absorbance spectra of probes **4**-**6** (10 μM) in the indicated formulations. **(D)** Quantification of fluorescent signal in cells by flow cytometry after incubation with probes formulated with and without addition of 1 % PEG-400. Mean value ± SD. n=3. Student’s t-test.

Another factor that could dramatically impact the efficiency of probe activation in cells is overall solubility which impacts the availability of the probes in aqueous buffers. To assess overall solubility, we took advantage of the fact that dye labeled molecules change both in intensity and excitation maxima depending on solubility and aggregation. This aggregation behavior has been described for cyanine dyes and likely further reduces availability in aqueous buffers [29]. We therefore measured absorption of the probes over a range of wavelengths near their absorption maximum of 650 nm. Reducing the amount of DMSO from 100% to 10% led to both a substantial drop of overall absorbance and a corresponding shift in the maximum signal to a peak at 600 nm, corresponding to aggregated probe (**Fig. 3C**). Furthermore, the biggest drop in absorbance upon reduction in DMSO levels was observed for probe **5** which has the most hydrophobic amino acid residues and contains a hydrophobic Cbz-capping group. To increase solubility, we added the *in vivo* compatible formulant polyethylene glycol-400 (PEG-400) to the solution. This substantially increased the solubility of all probes but was most effective for probe **5** and **6**, with both increasing to similar levels of overall absorption and peak shapes. We then tested the all three probes in cells using a 1:10 dilution of these solution in tissue culture media resulting in 1 μM probe with or without a final concentration of 1% PEG-400 (**Fig. 3D**). Interestingly, while the addition of PEG had similar positive effects on solubility of probes **5** and **6**, only probe **5** showed a 4-fold increase in signal upon PEG addition. This suggest that while poor solubility is a factor that limits the efficacy of probe **5**, the overall weak activity of probe **6** was not due to this issue, but likely due to ineffective uptake into cells.

### Tumor imaging with optimized probes

To test if the new probes, optimized based on the tumor proteolytic profile, yielded a higher fluorescent signal in tumors *in vivo*, we injected the probes in mice bearing 4T1 breast tumors. In confirmation of the insights gained from the cell labeling studies, we found that addition of 30% PEG-400 to our probe formulation strongly increases intensity of fluorescent signal *in vivo* (**Fig. S6**). Accounting for the dilution in the blood stream of the mouse, this amount of PEG-400 results in final concentration of about 0.7%. This enabled us to reduce the probe dose used in previous studies with 6QC (**4**) [14], by 5-fold from 2.3 mg/kg to 0.5 mg/kg for all three probes. The fluorescent signal in the tumor tissue was significantly higher using probe **5** compared to 6QC (**4**) (p=0.0007) or the acetylated version **6** (p=0.0002) (**Fig. 4A, B** and **Fig. S7**). This difference between probe signals is sufficient to be apparent by visual inspection, an important indicator that this is a relevant improvement for the use of these agents for intra operative surgical guidance (**Fig. 4A**). To quantitatively assess the signal enhancement of the new probes, **5** and **6**, we compared signals in the tumors to the adjacent mammary fat pad (**Fig. 4B**). As previously described, 6QC (**4**) effectively distinguished tumor from healthy fat pad, however the newly optimized probe **5** yielded a significantly higher (p<0.0001) signal in the tumor, while retaining overall low levels of signal in the normal, surrounding tissues resulting in a bright signal and higher signal over background with improved contrast (p=0.0007 for 6QC (**4**), p<0.0001 for probe **5**). The acetylated probe **6**, although most effectively processed in lysates, did not result in a significantly increased signal in tumors compared to background signal, likely due to its reduced cell permeability identified in our cellular studies (**Fig. 4B**). All of the substrate probes show fluorescent signal in the liver and kidneys, however these levels were not increased for probe **5** compared to the other probes, confirming that the increased activation within tumor is not simply due to overall increase availability of the probe (**Fig. 4C,D** and **Fig. S8**).

**Fig. 4:**
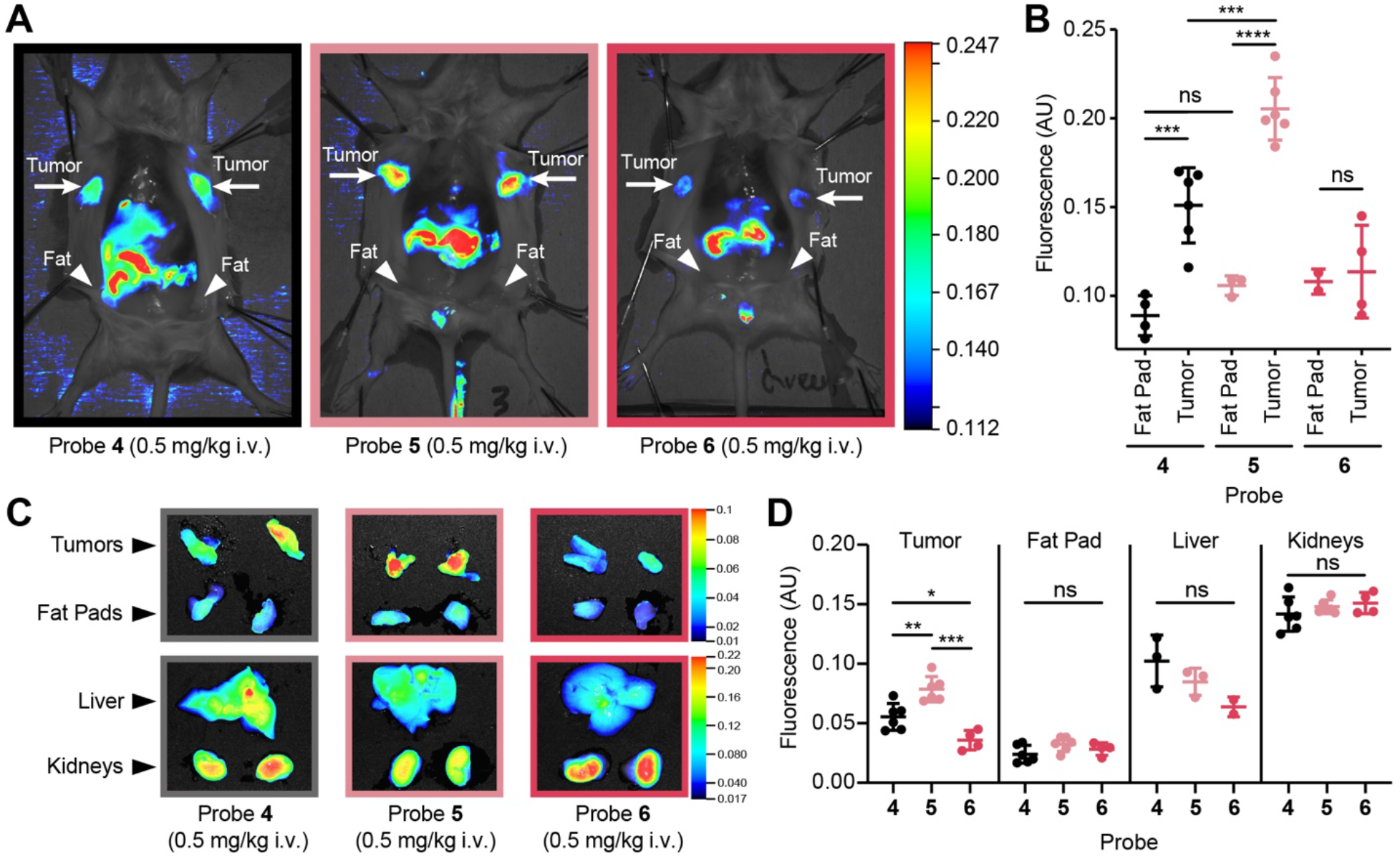
Imaging of breast tumors with optimized fluorescently quenched substrate probes. **(A)** Fluorescent signal in tumor bearing mice. Representative images of mice injected with probes **4**-**6** (0.5 mg/kg in 10% DMSO + 30% PEG-400, 3 h). Mice have been splayed to show mammary tumors. Location of tumors, and healthy fat pads are indicated **(B)** Quantification of fluorescence in healthy mammary fat pad and tumor tissue in 4T1 breast cancer mice injected with probes **4**-**6** (0.5 mg/kg in 10% DMSO + 30% PEG-400, 3 h, n_fat_= 3, n_tumor_= 6, Student’s t-test). **(C)** Representative *ex vivo* images of fluorescent signal in tumor, fat pad, liver, and kidneys for animals injected with probes **4**-**6**. **(D)** Quantification of ex vivo fluorescence in tumor, fat pad, liver, and kidneys in animals injected with probes **4**-**6** (n_tumor_= 6, n_fat_= 6, n_liver_= 3, n_kidney_= 6, Student’s t-test).

### Validation of optimized probes using human clinical samples

A major concern with an approach using mouse tumor tissues for probe optimization is that the resulting enhancements in the mouse will not translate into humans. To address this issue, we needed to evaluate if the overall patterns of activity of our probes in mouse tumor tissues were recapitulated in human specimens. We therefore applied all four of the optimized probes (**5**-**8**) and 6QC (**4**) to human tumor lysates derived from five breast cancer patients along with corresponding adjacent normal mammary tissue (histology shown in **Fig. 5A**). Quantification of the samples confirmed that, as observed in the mouse tissue, the acetylated substrates showed the highest cleavage rates, with substrate probe **6** being cleaved most efficiently by the clinical tumor lysates (**Fig. 5B** and **Fig. S9**). Importantly, when the proteolytic activity in the tumor samples was compared to adjacent normal breast tissue, all four optimized probes could robustly differentiate samples of cancerous tissue extracts from extracts derived from healthy tissues (p=0.0002 for **6**). In contrast, 6QC (**4**) was not efficiently cleaved in these lysates and therefore could not differentiate between cancer and normal tissue. Interestingly, the cleavage pattern of all five probes in human breast tumor lysates very closely resembled the proteolytic profile in mouse tumor tissue lysate and the preferences of cathepsin B (**Fig. 5C**).

**Fig. 5:**
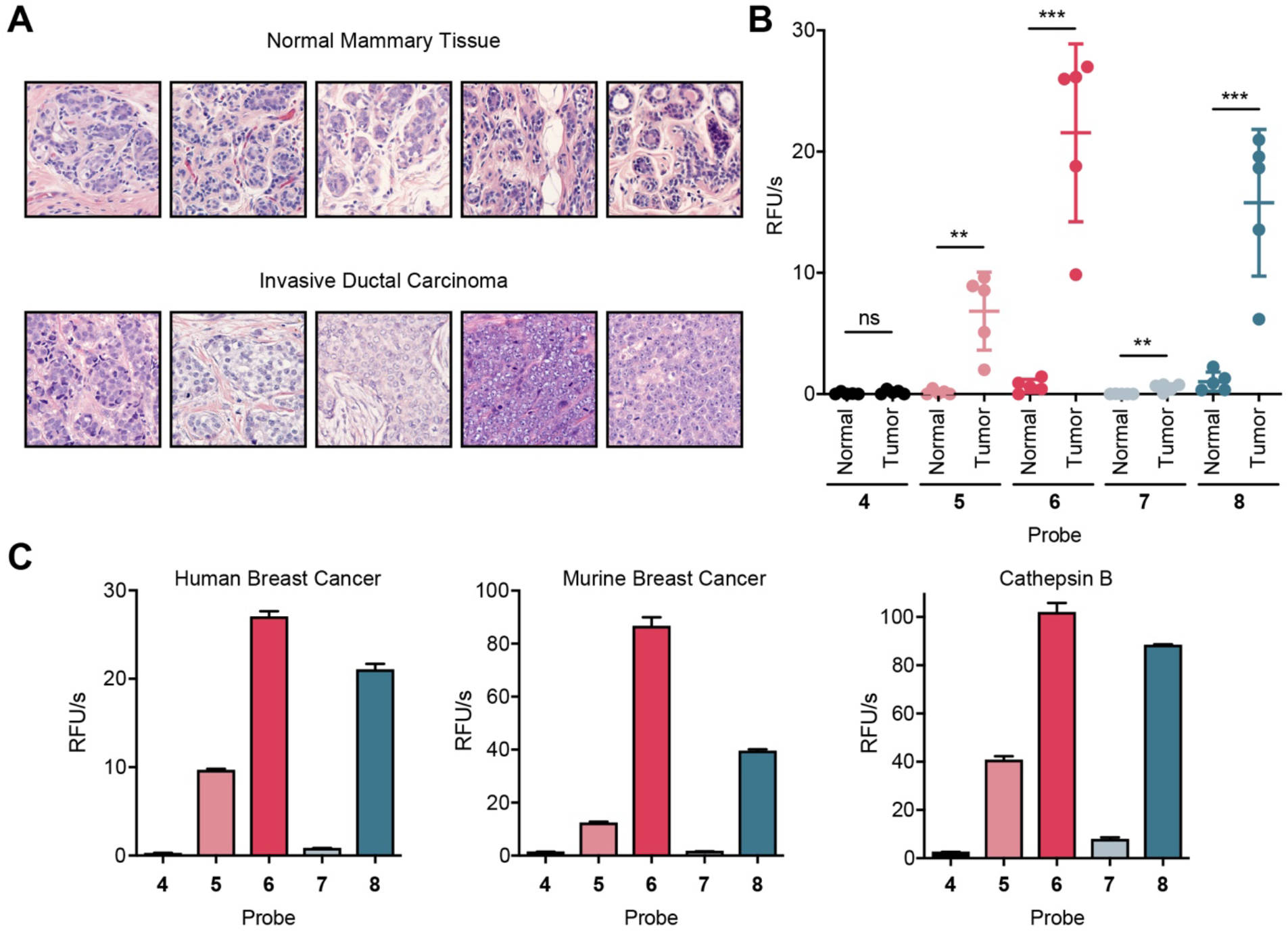
Optimized quenched substrate probes can distinguish human breast cancer samples. **(A)** Histology (Hematoxylin and eosin stain) of 5 patient biopsy samples of breast tumor tissue and adjacent healthy mammary tissue. **(B)** Quantification of cleavage rates of normal healthy human breast tissue (N) and human breast tumor tissue (T) incubated with **4-8** (n_Healthy_= 5, n_Tumor_= 5, Student’s t-test). **(C)** Comparison of cleavage pattern of one representative human breast cancer sample (left), the murine 4T1 tumor lysate (middle), and recombinant cathepsin B (right) incubated with probed **4-8**. Mean value ± SD. n=3.

## Discussion

Many of the current classes of probes being evaluated for fluorescence-based cancer imaging were designed based on a specific tumor expressed ligand or enzyme activity [4, 30]. This requires the selection of a target that may or may not be the most optimal ligand and then expending significant development efforts to generate a contrast agent. This can result in contrast agents that underperform or that could have been much more effective if other targets were prioritized. Therefore, we sought to develop methods to streamline the process of probe development while also allowing for an overall unbiased screening approach to identify optical reporters for the disease tissue of interest. We therefore performed screens for signatures of protease activity directly in diseased tissue extracts. This allows rapid identification of lead scaffolds that have been selected based solely on their ability to be cleaved in the disease tissue of interest. This approach does not depend on knowledge of a target or cell type of origin and probes designed based on tissue screening data can be quickly evaluated in animal models with less iterative synthetic chemical optimization steps.

In this study, we focused on mouse breast cancer tissue lysate to determine optimal peptide sequences to incorporate into the design of the probes. The resulting probes showed increased overall signals in tumor tissues compared to our previously reported probe, 6QC (probe 4), that was designed to target cathepsins. Interestingly, we found that the optimal probes from the tissue profiling studies were predominantly processed by cathepsin B. This allowed us to compare the selectivity of our new probes to existing specificity data reported for purified cathepsins that had been screened using the same substrate library [25]. This analysis confirmed that top amino acids that we selected from the lysate screens matched the optimal residues for the purified cathepsin B screen. However, the published study using purified cathepsins was focused on designing probes selective for individual cathepsins. As a result, other, less optimal residues were chosen to produce probes. This difference nicely highlights the value of using the unbiased approach presented here for the design of in vivo optical probes, as it focuses on the selection of the most effective probe regardless of target.

In order to gain further tumor tissue selectivity, we could also perform counter screening of the substrate library with extracts derived from normal ‘healthy’ tissues. However, it can often be difficult to obtain a homogeneous sample that is an accurate representation of surrounding healthy tissues. Furthermore, such normal tissues must have a clear fingerprint of activity that could be used to negatively select for residues. We have performed screens of normal kidney tissue, given its high level of signal for most probes (data not shown). However, we found that the same enzyme signatures (i.e. cathepsins) dominated the profile and therefore did not provide useful data for negative selection. Regardless. We show here that by focusing on the substrates with the most optimal activity in the disease tissue, it is possible to generate a probe with enhanced signal in those tissues compared to healthy organs such as kidney, that also express many of the same enzymes.

Our optimized probes were not only brighter than previously, classically designed probes but were able to distinguish human breast cancer from adjacent mammary tissue. It is notable that the probe cleavage pattern found in mouse breast cancer tissues is remarkably similar to that of human breast cancer tissue. This suggests that tissue from relevant animal models may serve as an accurate representation of the proteolytic signature of human disease. This finding for breast cancer tissues is consistent with the fact that many proteases have highly conserved physiological and pathological functions (i.e. blood coagulation, tissue remodeling in inflammation and wound healing) across diverse organisms. These conserved roles often lead to conserved specificity, making animal models a reasonable surrogate for human disease when assessing proteolytic signatures.

Applying this tissue screening approach, we identified an optimized substrate probe that was about 65 times more efficiently cleaved than our previous probe 6QC (**4**) in *in vitro* assays. However, our results show that to optimize these probes for *in vivo* applications, other properties must be carefully considered. We found that an optimal next step in probe optimization is to assess labeling efficiency in cellular culture systems. In particular, quantitative flow cytometry analysis proved to be a valuable tool to accurately assess the translatability from *in vitro* assays to *in vivo* mouse studies. In this assay, we observed that properties of the probe other than the peptide sequence greatly influenced cell labeling. N-terminal capping with a bulky aromatic carboxybenzyl group increased cell labeling compared to molecules containing a smaller acetyl group. All of our results suggest this increase in due to improved cellular uptake. Furthermore, we used the flow cytometry analysis to determine how formulating the probe to improve solubility impacted cell labeling. We found the most hydrophobic scaffold (**5**) greatly benefited from addition of PEG-400 as a formulant. Our results from the *in vivo* imaging studies confirmed that use of optimal formulation increase fluorescent signals in tumors and therefore allowed us to reduce probe dosage by as much as 5-fold while still retaining optimal tumor contrast.

Our biodistribution studies with our new set of probes (**5**-**8**) also allowed us to evaluate basic PK/PD properties of these substrate probes. Small molecule peptide cathepsin probes tend to accumulate in the liver and kidneys due to clearance mechanisms for these types of agents. The relatively high physiological cathepsin activity in these organs results in large background signals from most probes. However, in contrast to the tumor signal, our optimized probe (**5**) did not show an increase in signal in the liver and kidneys compared to the 6QC (**4**) probe. Furthermore, the acetylated derivative of the optimized probe (**6**) showed almost no signal accumulation in the tumors, but had the same high signal intensity in the kidney as the 6QC (**4**) probe. This suggests that the observed background signals of all probes in the liver and kidney are likely due to increased probe uptake and exposure whereas the tumor contrast reflects the optimal substrate processing observed in the tumor tissue extracts.

In this study, we used human tumor tissues for profiling using the optimized imaging probes. We chose to use human breast tumor tissues because our screening data was performed in a mouse model of breast cancer. This analysis of human tissues was enabled by the relatively easy access of breast tissues biopsies of significant size to enable profiling. However, a limited amount of tissue may pose challenges for some disease indications (i.e. altherosclortic plaques, micro metastases, etc). In our current studies we performed screening of libraries in 96 well format plates using significant volumes (μLs), which required substantial amounts of tumor lysates (approximately 2 mg total protein per 96 well plate). However, the approach for screening for substrate processing is amenable to microwell plates (i.e. >384 wells) as well as suitable for adaptation to a coded bead-based HTS platform. In fact, Bachovchin et al. have described methods for bead-based readout of fluorescent probe competition assay to profile selectivity of compounds against diverse purified enzymes [31]. The same approach could allow rapid screening of sample-limited disease tissues for activity against many diverse enzyme substrate libraries. In principle, this approach should work for identification of optimal binding molecules or substrates for any enzyme for which an on-bead assay can be developed. By miniaturizing the screening assay, it should be possible to apply this approach to many diverse types of disease states, including liquid biopsies.

In summary, we demonstrate how screening a crude disease tissue lysate with diverse peptide substrate libraries can streamline the development of small molecule optical contrasts agents. It demonstrates the value of an application-oriented approach to probe design by ensuring efficient cleavage in the relevant tissue of interest as the first step in development. We believe this strategy will be broadly applicable to other diseases for which optical contrast agents could benefit disease management.

## Supporting information

Tholen et al Supporting Information

## Contributions

M.T., J.J.Y., and M.B. designed the study. M.T. performed *in vitro* experiments. J.J.Y. and E.Y. synthesized compounds. M.T. and J.J.Y. performed *in vivo* experiments. MD contributed combinatorial HyCoSuL libraries. K.G. ran samples on libraries. B.A.M. and N.S.vdB. helped with human sample experiments. M.T., J.J.Y. and M.B. wrote the text. All authors reviewed text.

## Notes

The authors do not declare any conflict of interests.

## Acknowledgements

This work was supported by NIH Grant R01 EB026285 (to M.B.) and a Stanford Cancer Institute Translational Oncology Program seed grant (to M.B.), DFG Research Fellowship TH2139/1-1 (to M.T.), and Stanford ChEM-H Chemistry/Biology Interface Predoctoral Training Program and NSF Graduate Research Fellowship Grant DGE-114747 (to J.J.Y.). Marcin Drag laboratory is supported by the “TEAM/2017-4/32” project, which is conducted within the TEAM programme of the Foundation for Polish Science cofinanced by the European Union under the European Regional Development Fund and National Science Centre in Poland. The authors would like to thank the Turk Lab at J. Stefan Institute for providing the recombinant cathepsin proteases used in this study. The authors also thank the members of the Bogyo Group for helpful discussions.

