## Supplementary material for "Design of optical imaging probes by screening of diverse substrate libraries directly in disease tissue extracts": Tholen et al Supporting Information

**Table of Contents**


1. Supplementary Materials and Methods S3

2. Chemical Synthesis and Characterization
   - Synthetic Scheme of Compounds 1-3 S4
   - General Synthetic Procedure for Compounds 1-3 S5
   - LCMS traces of Compounds 1-3 S5-6
   - Synthesis Scheme of Compounds 5-8 S7
   - General Synthetic Procedure for Compounds 5-8 S8
   - LCMS traces of Compounds 5-8 S9-12

3. Supporting Figures S13-20

4. References S21

### 1. Supplementary Materials and Methods

All reagents were purchased from commercial suppliers and used without further purification. Sulfo-Cy5 NHS ester and QSY-21 NHS ester were purchased from commercial suppliers. Reagent grade solvents dimethylformamide (DMF) and dichloromethane (DCM) were used for chemical reactions. Reactions were analyzed by LC-MS using an API 150EX single-quadrupole mass spectrometer (Applied Biosystems). Synthesized compounds were purified via reverse-phase HPLC using a 1260 Infinity II LC System (Agilent Technologies) equipped with a C18 column. Column used was a Luna® 5 µm C18(2) 100 Å LC Column 250 mm x 10 mm (Phenomenex). Compounds were eluted with a gradient of distilled water and acetonitrile containing 0.1 % trifluoracetic acid as solvents.

Abbreviations: DIPEA = N,N-Diisopropylethylamine, TFA = trifluoroacetic acid

### 2. Chemical Synthesis and Characterization

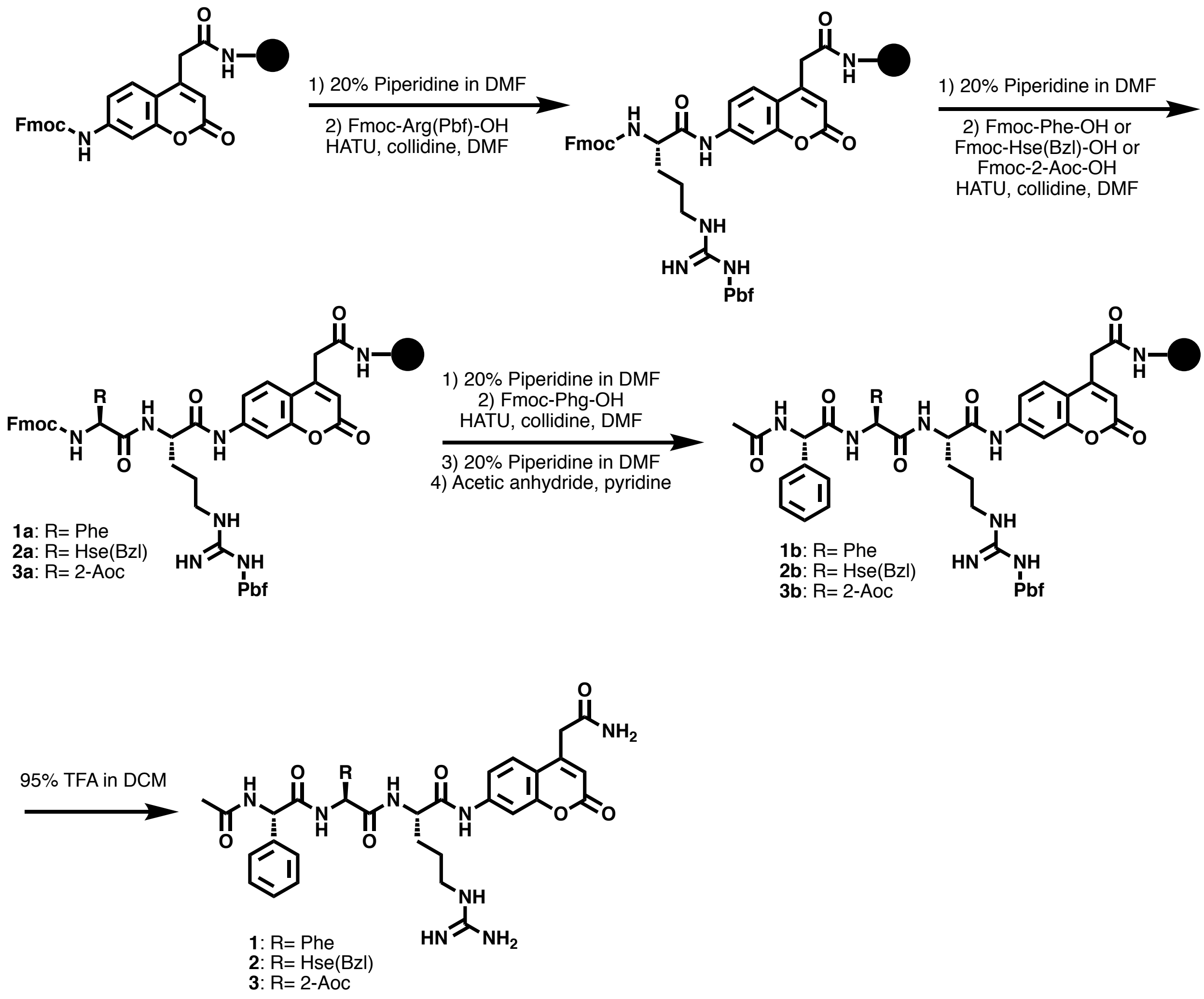


**Scheme S1.** Synthesis of compounds **1-3** using combination of solid and solution phase synthesis.

General synthetic procedure for peptides **1-3**

1. Swell ACC-resin[1] with DMF for 30 min in Solid Phase Peptide Synthesis (SPPS) vessel.
2. Deprotect Fmoc group with 20% piperidine in DMF for 30 min.
3. Drain and wash with DMF, add Fmoc-Arg(Pbf)-OH (5 eq.), HATU (5 eq.), collidine (5 eq) in DMF and shake O/N.
4. Drain and wash with DMF and carry out additional coupling with Fmoc-Arg(Pbf)-OH (5 eq.), HATU (5 eq.), collidine (5 eq) in DMF and shake O/N.
5. Drain and wash with DMF and acetylate unreacted ACC with 20% $Ac_2O$ in pyridine for 1 h.
6. Drain and wash with DMF 5 times and with DCM 5 times.
7. Drain and wash with DMF and deprotect Fmoc group by adding 20% piperidine in DMF and shake for 30 min.
8. Drain and wash with DMF and add Fmoc-X-OH (X = Phe, Hse(Bzl), or Fmoc-2-Aoc-OH, 3.0 eq.), HATU (5 eq.), collidine (5 eq) in DMF (8 mL) and shake for 1 h.
9. Drain and wash with DMF and deprotect Fmoc group by adding 20% piperidine in DMF and shake for 30 min.
10. Drain and wash with DMF and add Fmoc-Phg-OH (3.0 eq.), HATU (5 eq.), collidine (5 eq) in DMF (8 mL) and shake for 1 h.
11. Drain and wash with DMF and deprotect Fmoc group by adding 20% piperidine in DMF and shake for 30 min.
12. Drain and wash with DMF and cap peptide with 20% $Ac_2O$ in pyridine for 1 h.
13. Cleave peptide off resin with 95% TFA in DCM and collect wash. Evaporate to dryness.

Compound **1** LCMS Trace

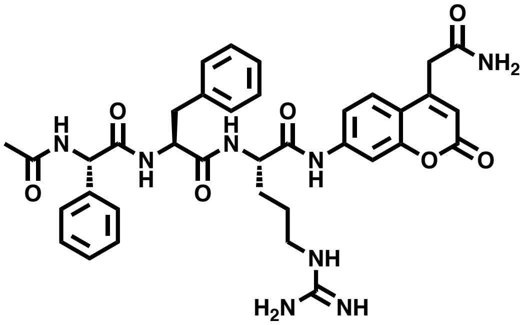

**Compound 1**

Chemical Formula: $C_{36}H_{40}N_8O_7$
Exact Mass: 696.30
Molecular Weight: 696.77

ESI-MS (m/z) for $C_{36}H_{41}N_8O_7^{1+}$
Calculated: 697.3
Found 697.4

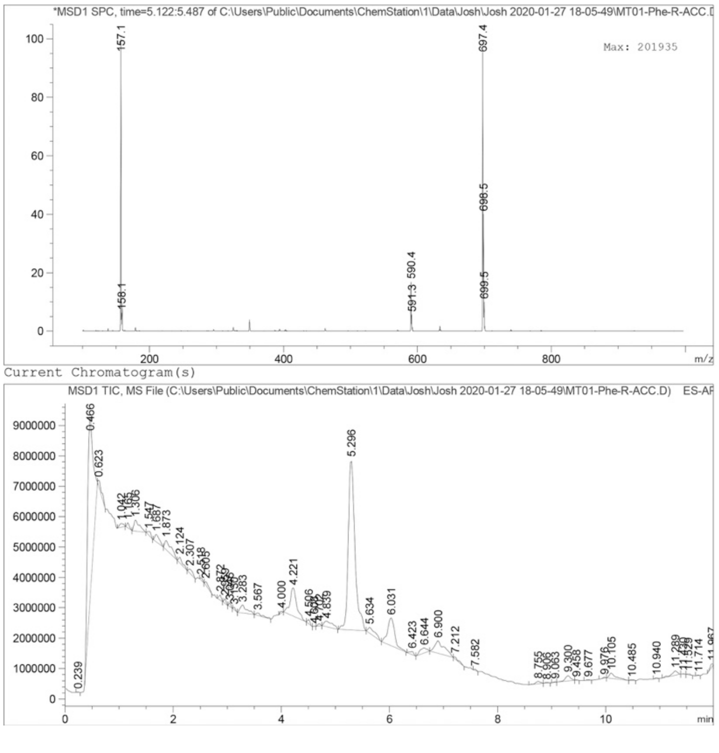

Compound **2** LCMS Trace

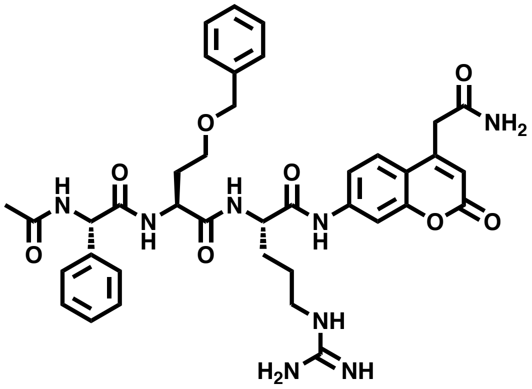

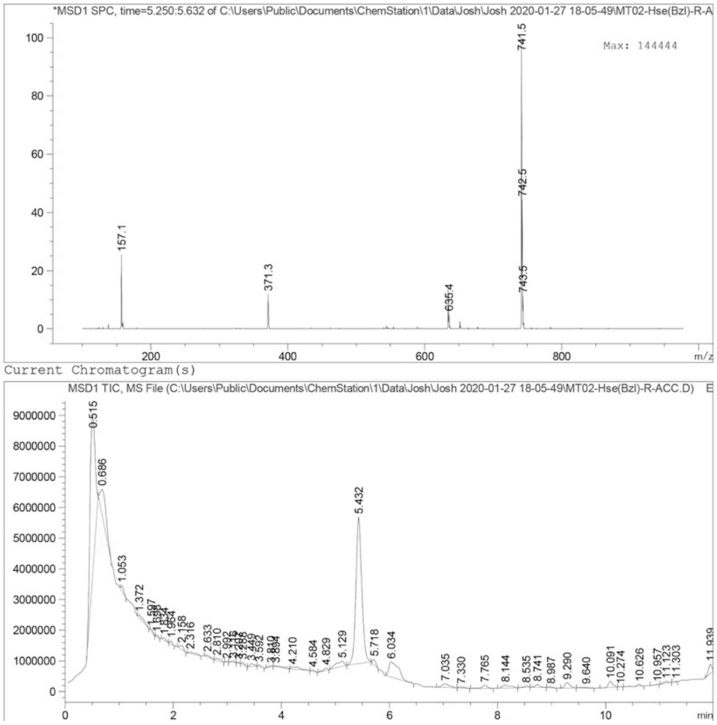


**Compound 2**

Chemical Formula: $C_{38}H_{44}N_8O_8$
Exact Mass: 740.33
Molecular Weight: 740.82

ESI-MS (m/z) for $C_{38}H_{45}N_8O_8^{1+}$
Calculated: 741.3
Found 741.5

Compound **3** LCMS Trace

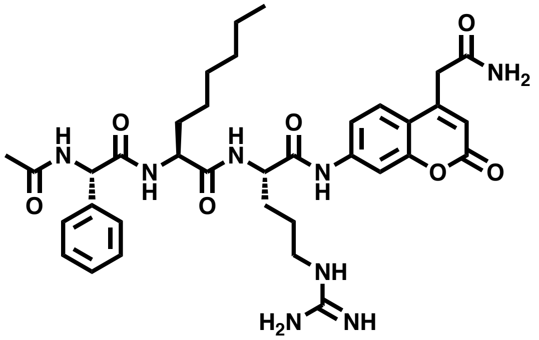

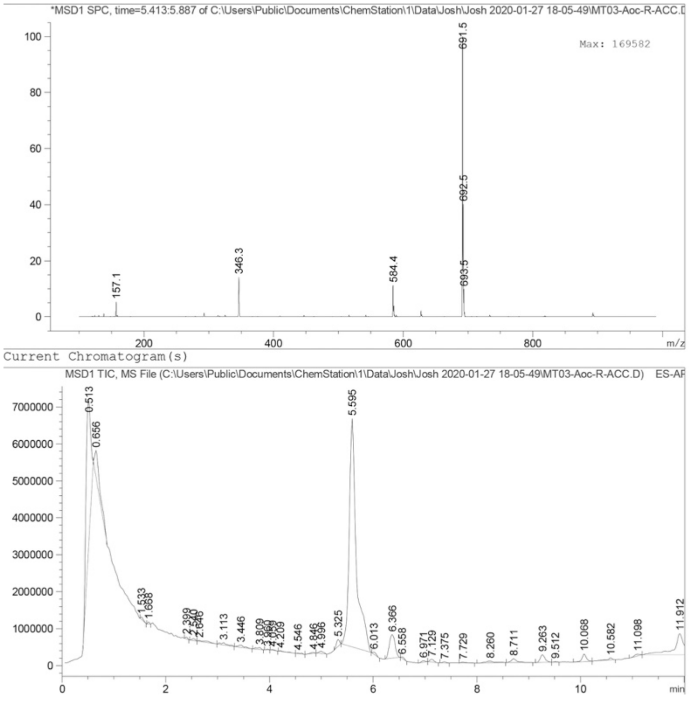


**Compound 3**

Chemical Formula: $C_{35}H_{46}N_8O_7$
Exact Mass: 690.35
Molecular Weight: 690.80

ESI-MS (m/z) for $C_{35}H_{47}N_8O_7^{1+}$
Calculated: 691.4
Found 691.5

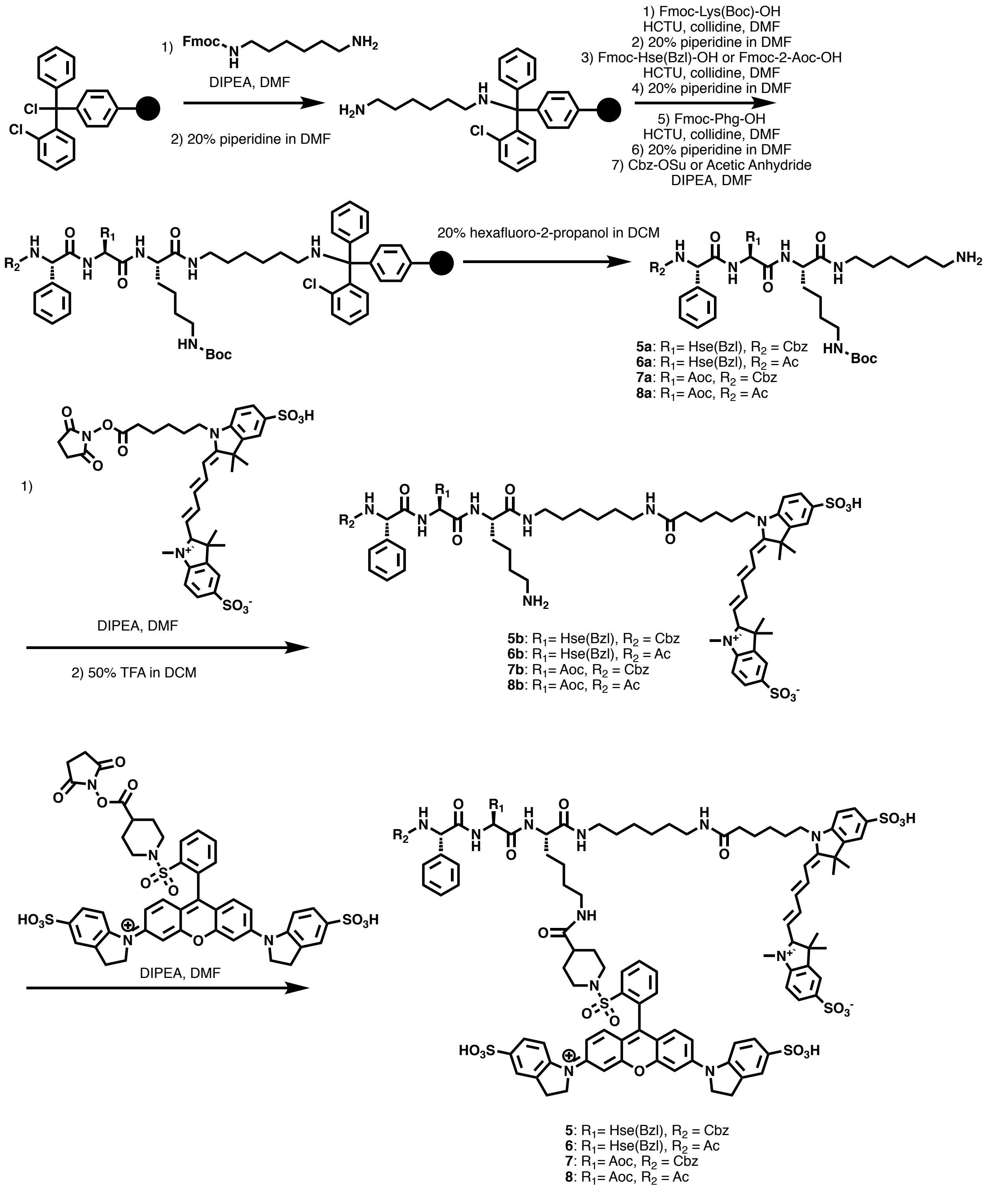


**Scheme S2.** Synthesis of compounds **5-8** using combination of solid and solution phase synthesis.

General synthetic procedure[2] for peptides **5a-8a**

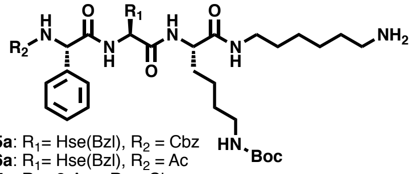

**5a**: $R_1$= Hse(Bzl), $R_2$ = Cbz
**6a**: $R_1$= Hse(Bzl), $R_2$ = Ac
**7a**: $R_1$= 2-Aoc, $R_2$ = Cbz
**8a**: $R_1$= 2-Aoc, $R_2$ = Ac

1. Weigh ~200mg resin in Solid Phase peptide synthesis (SPPS) vessel and swell resin with DCM for 30 min.
2. Drain DCM and add mono-Fmoc-diaminohexane (1.5 eq. of resin), DIPEA (8.0 eq.), DMF (8 mL) and shake O/N.
3. Drain and wash resin with methanol and add methanol and shake for 30 min.
4. Drain and wash with DMF and add 20% piperidine in DMF and shake for 30 min.
5. Drain and wash with DMF and add Fmoc-Lys(Boc)-OH (3.0 eq.), HCTU (4.5 eq.), collidine (4.5 eq) in DMF (8mL) and shake for 1 h.
6. Drain and wash with DMF and add 20% piperidine in DMF and shake for 30 min.
7. Drain and wash with DMF and add Fmoc-X-OH (X = Hse(Bzl) or Fmoc-2-Aoc-OH, 3.0 eq.), HCTU (4.5 eq.), collidine (4.5 eq) in DMF (8mL) and shake for 1 h.
8. Drain and wash with DMF and add 20% piperidine in DMF and shake for 30 min.
9. Drain and wash with DMF and add Fmoc-Phg-OH (3.0 eq.), HCTU (4.5 eq.), collidine (4.5 eq) in DMF (8mL) and shake for 1 h.
10. Drain and wash with DMF and add 20% piperidine in DMF and shake for 30 min.
11. Drain and wash with DMF and add Cbz-Osu (10 eq.) or acetic anhydride (10 eq.), DIPEA (20 eq.), in DMF (8mL) and shake for 1 h.
12. Cleave peptide with 20% hexafluoro-2-propanol in DCM and collect wash. Evaporate to dryness.
13. Reverse-phase HPLC purification of peptides resuspended in 50% acetonitrile in water with 0.1% TFA.

General synthetic procedure for probes **5-8**

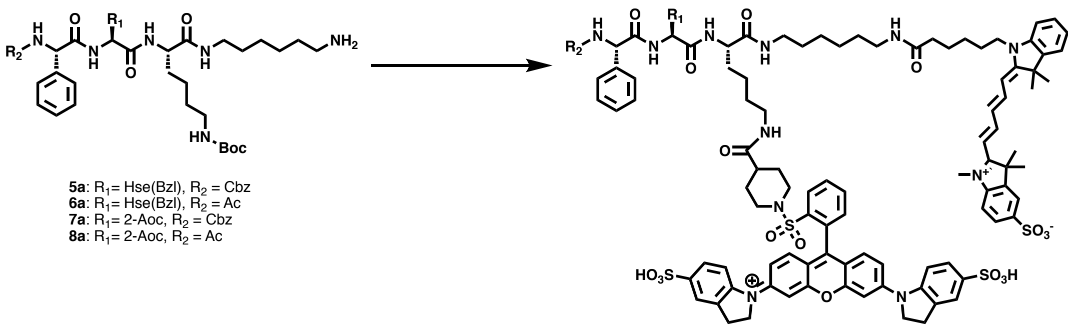

**5a**: $R_1$= Hse(Bzl), $R_2$ = Cbz
**6a**: $R_1$= Hse(Bzl), $R_2$ = Ac
**7a**: $R_1$= 2-Aoc, $R_2$ = Cbz
**8a**: $R_1$= 2-Aoc, $R_2$ = Ac

**5**: $R_1$= Hse(Bzl), $R_2$ = Cbz
**6**: $R_1$= Hse(Bzl), $R_2$ = Ac
**7**: $R_1$= 2-Aoc, $R_2$ = Cbz
**8**: $R_1$= 2-Aoc, $R_2$ = Ac

To the starting peptide was added sulfo-Cy5-NHS ester (1.5 eq.) + DIPEA (10 eq.) in DMF (400 uL) in 1.5 mL Eppendorf tube (covered from light) and shook for 3 h at room temperature. Reaction mixture was purified via HPLC and product was lyophilized. The powder was subjected to 50% TFA in DCM (1 mL) for 2 h and then the solvent was evaporate in vacuo. To the dry compound was added sulfo-QSY-21 NHS ester (1.5 eq.) + DIPEA (10 eq.) in DMF (400 uL) in 1.5 mL Eppendorf tube (covered from light) and shook for 3 h at room temperature. Reaction mixture was purified via HPLC and product was lyophilized and kept at -20 °C.

Compound **5** LCMS Trace

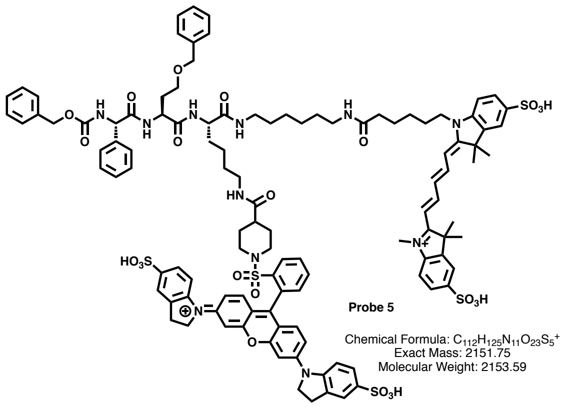


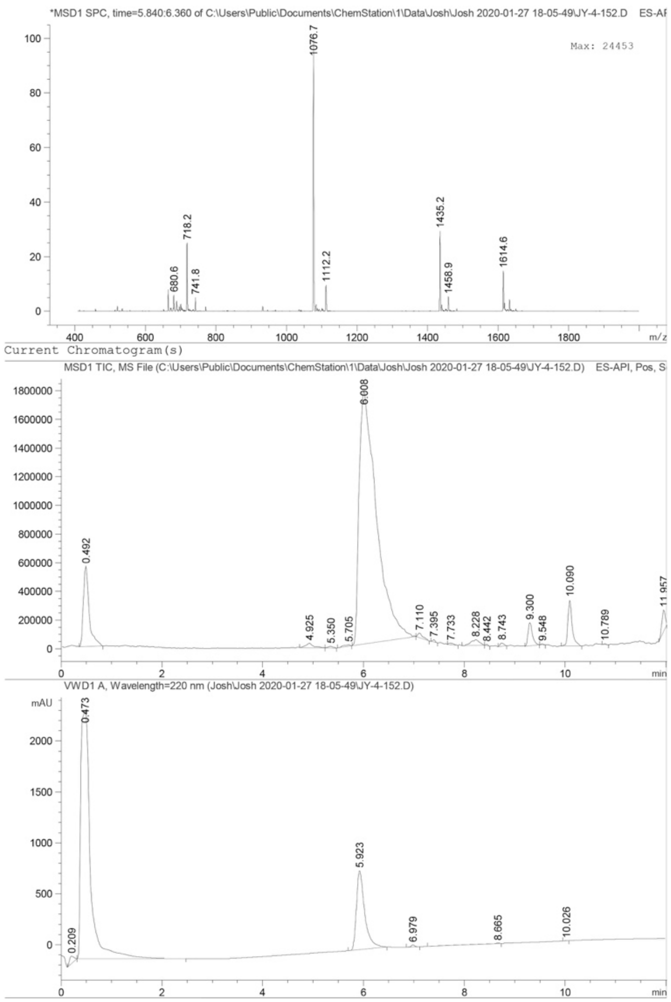


ESI-MS (m/z) for $C_{112}H_{126}N_{11}O_{23}S_5^{2+}$
Calculated: 1076.9
Found 1076.7

Compound **6** LCMS Trace

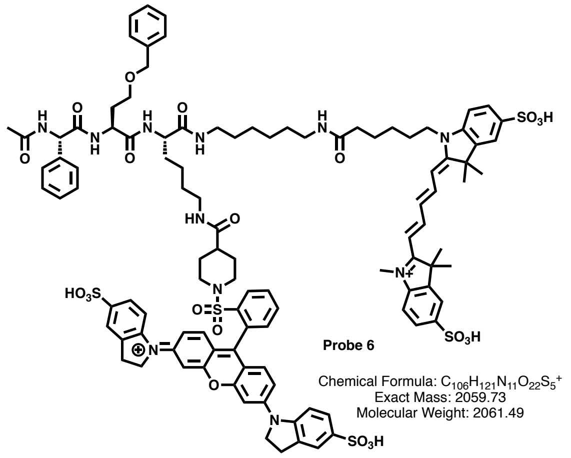


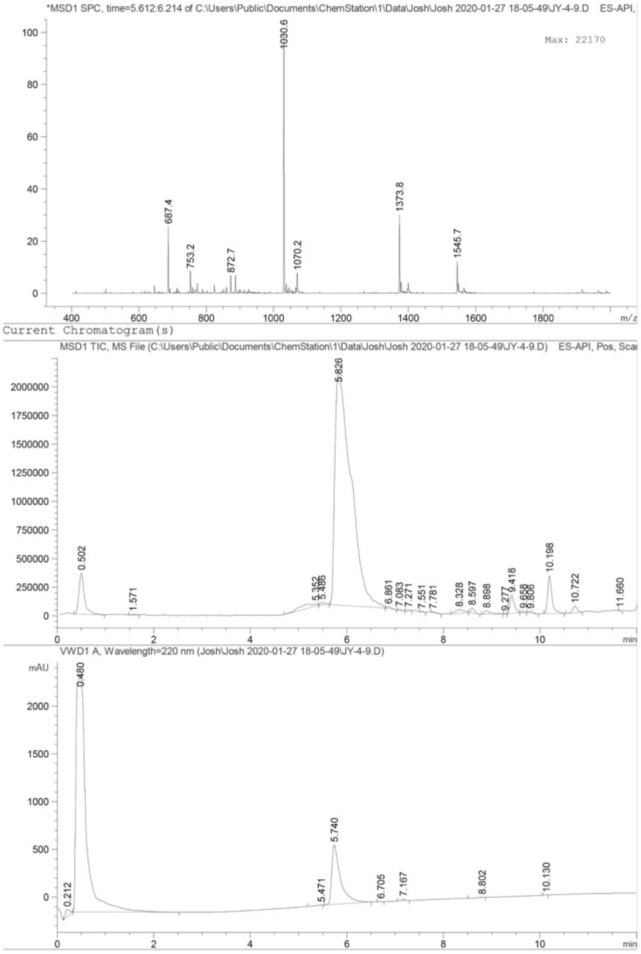


ESI-MS (m/z) for $C_{106}H_{122}N_{11}O_{22}S_5^{2+}$
Calculated: 1030.9
Found 1030.6

Compound **7** LCMS Trace

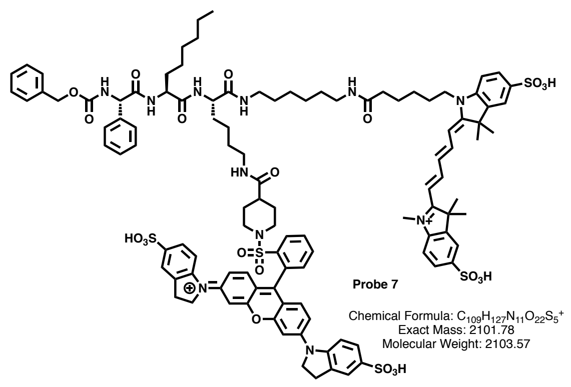


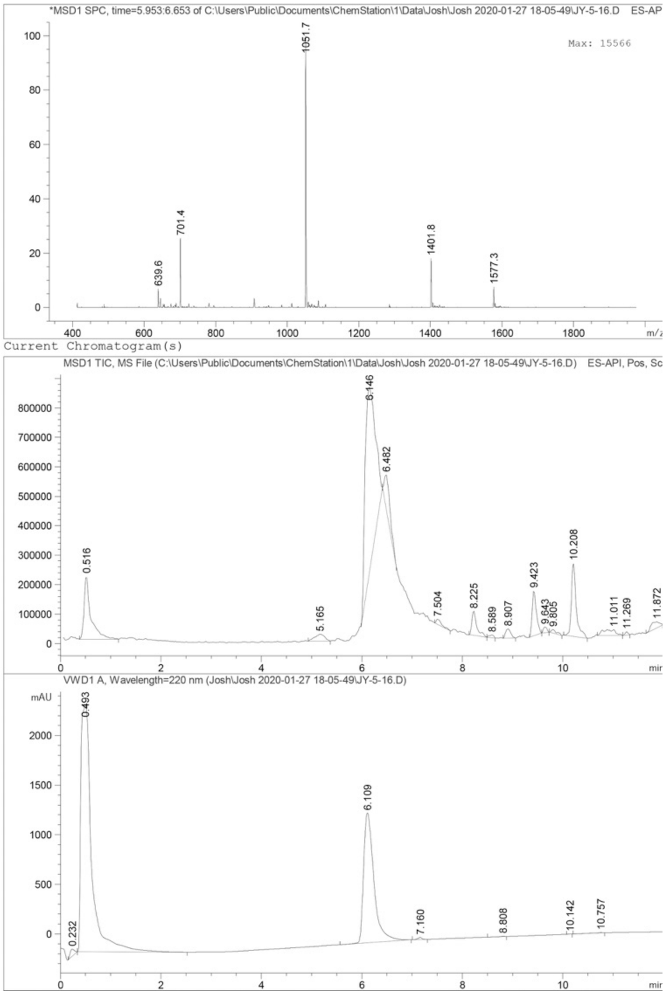

ESI-MS (m/z) for $C_{109}H_{128}N_{11}O_{22}S_5^{2+}$
Calculated: 1051.9
Found 1051.7

Compound **8** LCMS Trace

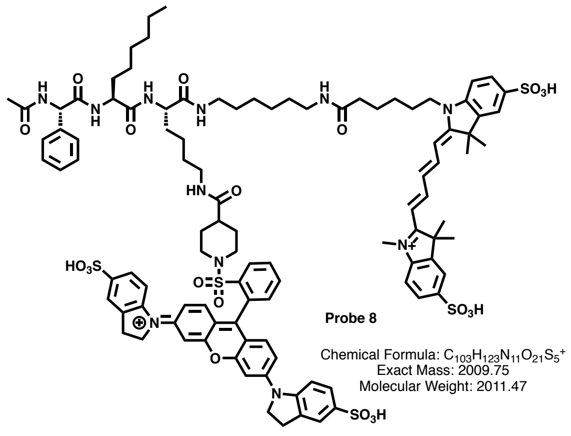


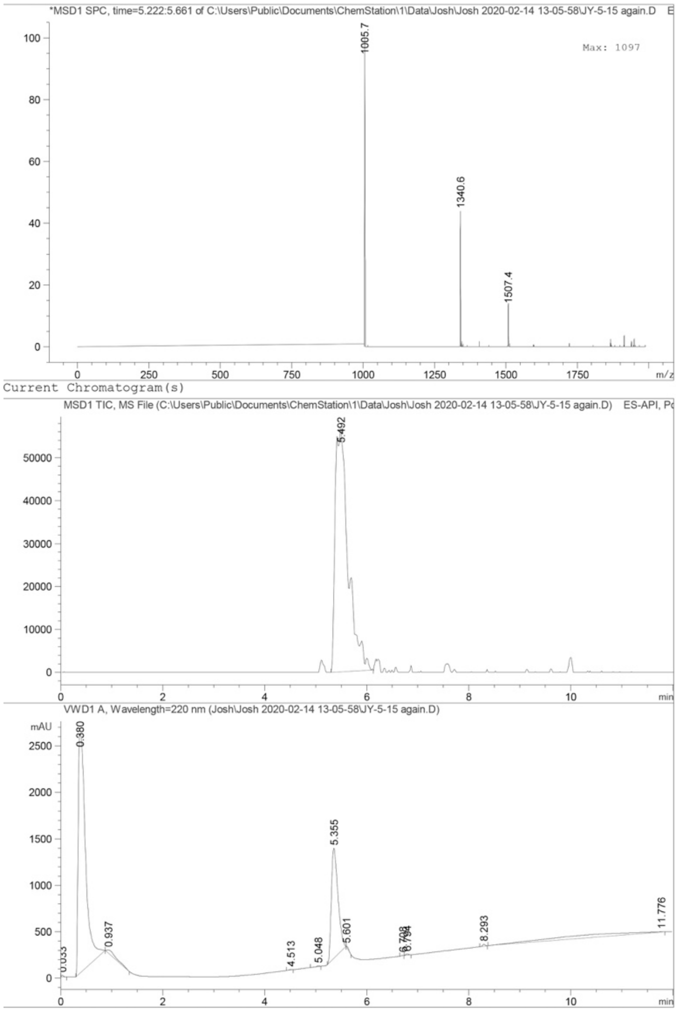


ESI-MS (m/z) for $C_{103}H_{124}N_{11}O_{21}S_5^{2+}$
Calculated: 1005.9
Found: 1005.7

### 3. Supporting Figures

A

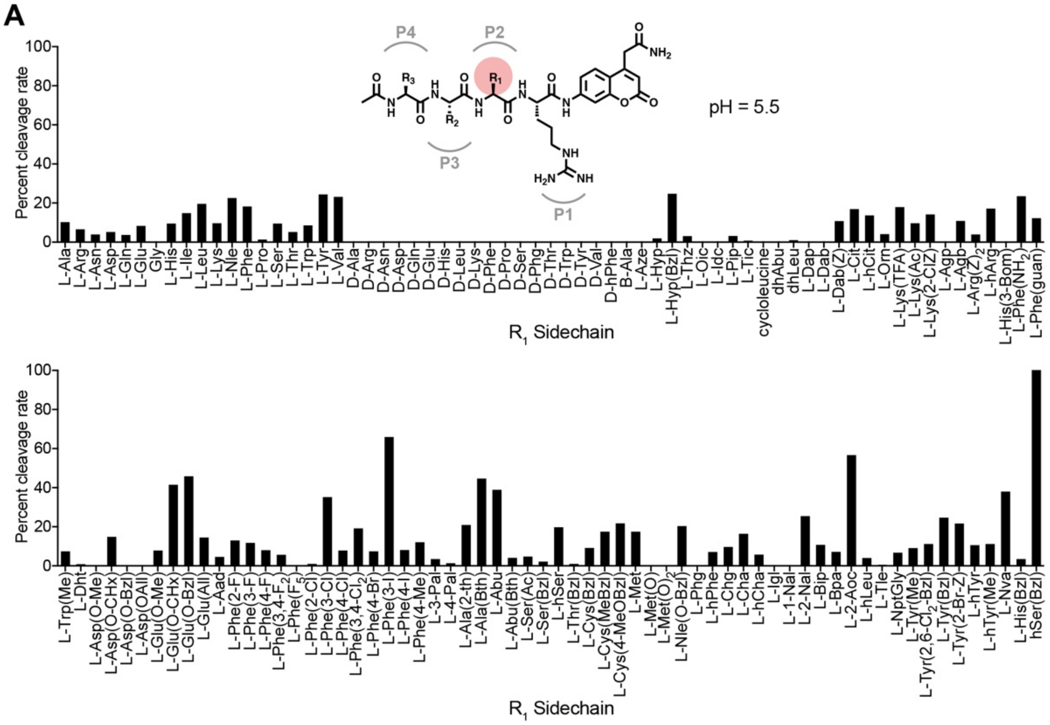


B

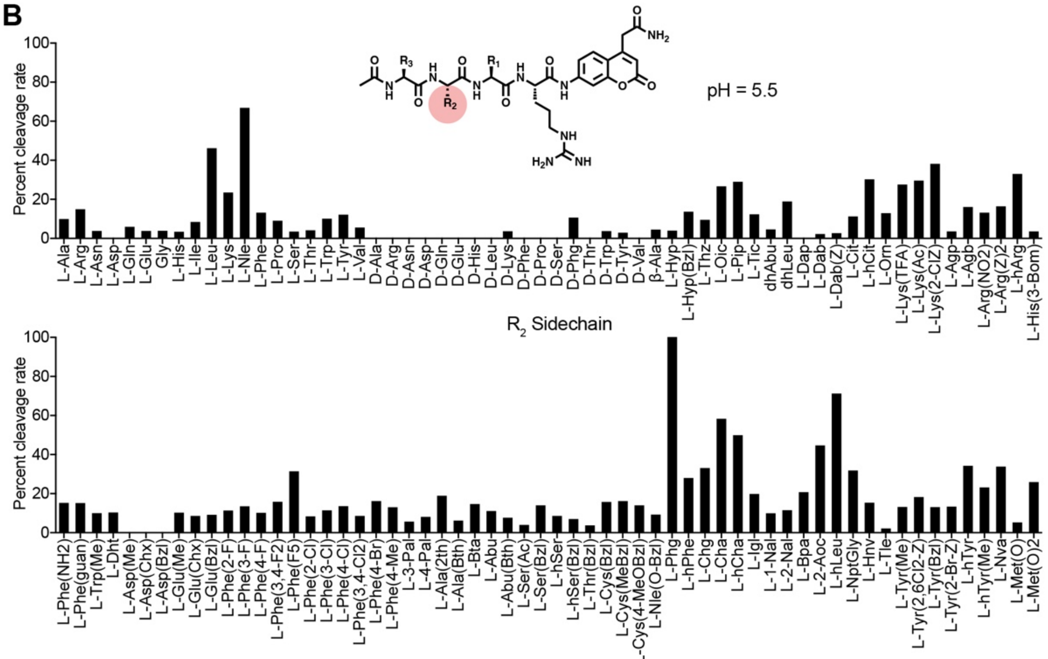


**Fig. S1: Relative cleavage rates in tumor lysate for all substrates in the HyCoSul library at pH 5.5.** **(A)** Screening of the P2 scanning HyCoSuL library in mouse tumor tissue extract. Plot of ACC cleavage relative to the most effectively cleaved residue for the full set of substrates at pH 5.5. Mean values of two tumor measurements were used. **(B)** Screening of the P3 scanning at pH 5.5. The P3 scanning library was assayed exactly as in (A).

**A**

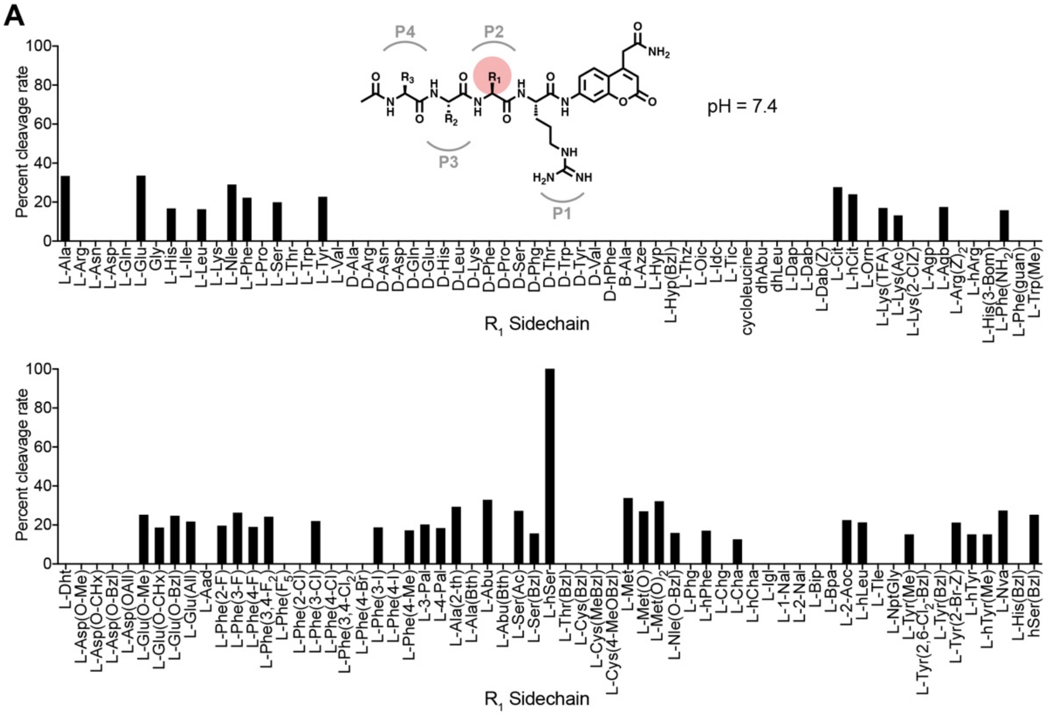


**B**

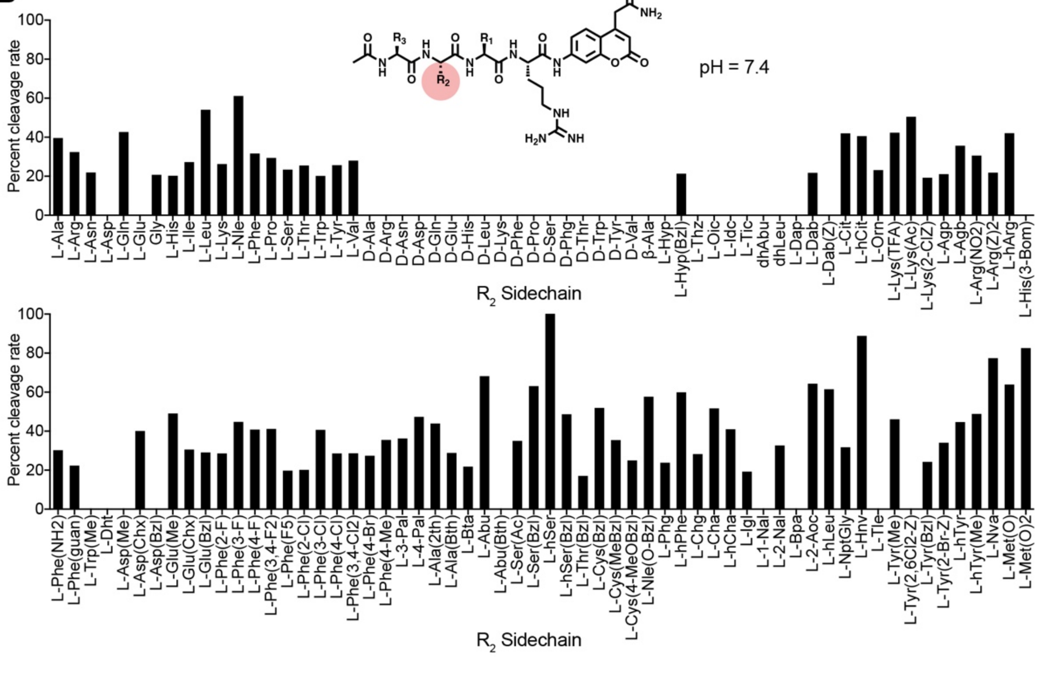


**Fig. S2: Relative cleavage rates in tumor lysate for all substrates in the HyCoSul library at pH 7.4.** **(A)** Screening of the P2 scanning HyCoSuL library in mouse tumor tissue extract. Plot of ACC cleavage relative to the most effectively cleaved residue for the full set of substrates at pH 7.4. Mean values of two tumor measurements were used. **(B)** Screening of the P3 scanning at pH 7.4. The P3 scanning library was assayed exactly as in (A).

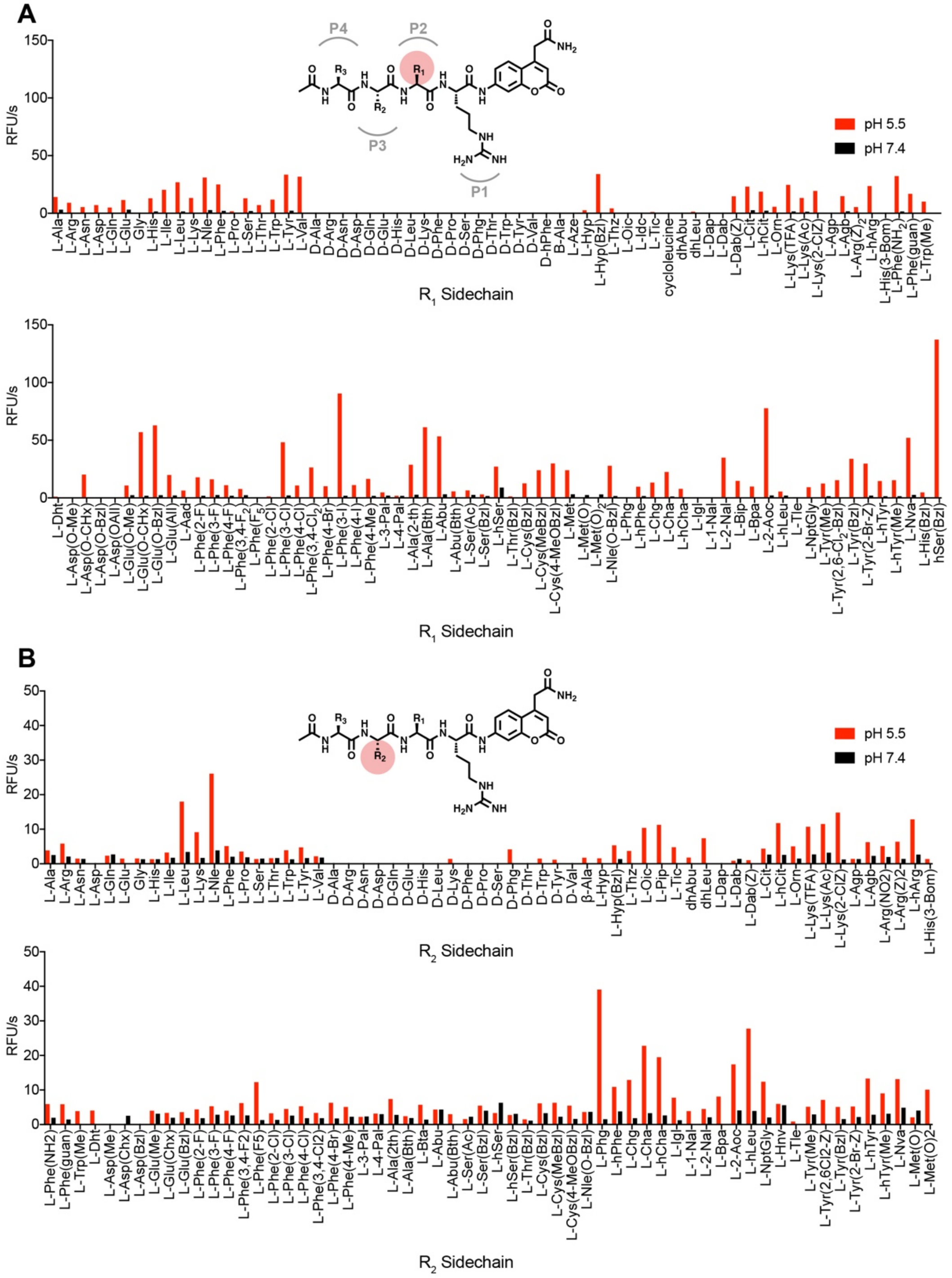


**Fig. S3: Comparison of raw cleavage rates at pH 5.5 and pH 7.4.**
**(A)** Screening of the P2 scanning HyCoSuL library in mouse tumor tissue extract. Plot of raw ACC cleavage rates at pH 5.5 (red) and pH 7.4 (black) for the full set of substrates. Mean values of two tumor measurements were used. **(B)** Comparison of the screening rates for the P3 scanning.

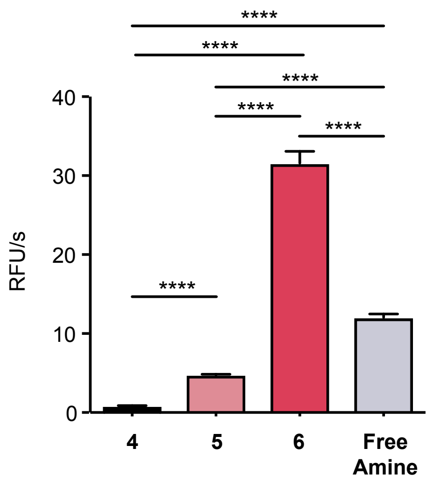


**Fig. S4: Cleavage rates for fluorescently quenched substrate probes in lysate of immortalized macrophages.**
Cleavage rate of probes **4-6** and the free amine version of **6** in lysate of immortalized macrophages at pH 5.5. Values are plotted as mean value ± SD. n=3.

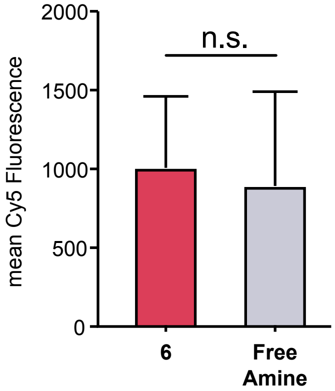


**Fig. S5: Comparison of in cell labelling with acetylated or uncapped version of optimized fluorescently quenched substrate probe 6.**
Quantification of fluorescent signals in immortalized macrophages incubated with probe **6** and an uncapped version of **6** with an N-terminal free amine (3 h incubation, 10 µM compound). Mean value ± SD. n=3. Student's t-test.

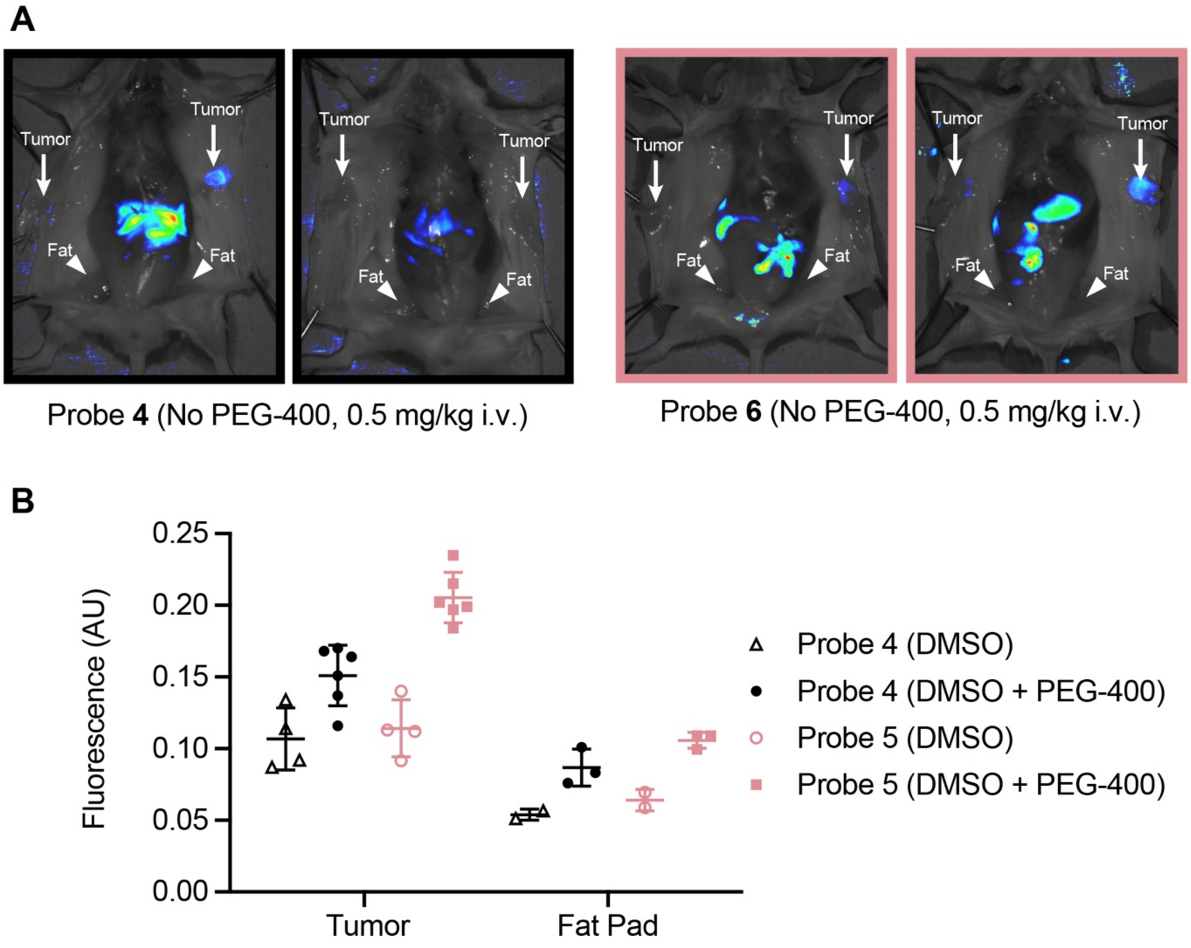


**Fig. S6: Effect of PEG-400 formulation of probes *in vivo*.**
**(A)** Fluorescent signal in tumor bearing mice. Images of mice injected with 0.5 mg/kg of probes **4** and **5** either in 10% DMSO / PBS or 10% DMSO / 30% PEG-400 / PBS, 3 h. Mice have been splayed to show mammary tumors. Location of tumors, and healthy fat pads are indicated. (**B**) Quantification of fluorescence in healthy mammary fat pad and tumor tissue in 4T1 breast cancer mice injected with 0.5 mg/kg of probes **4** and **5** with or without the PEG-containing formulation shown in (A), 3 h, $n_{fat}$= 2, $n_{tumor}$= 4, Student's t-test.

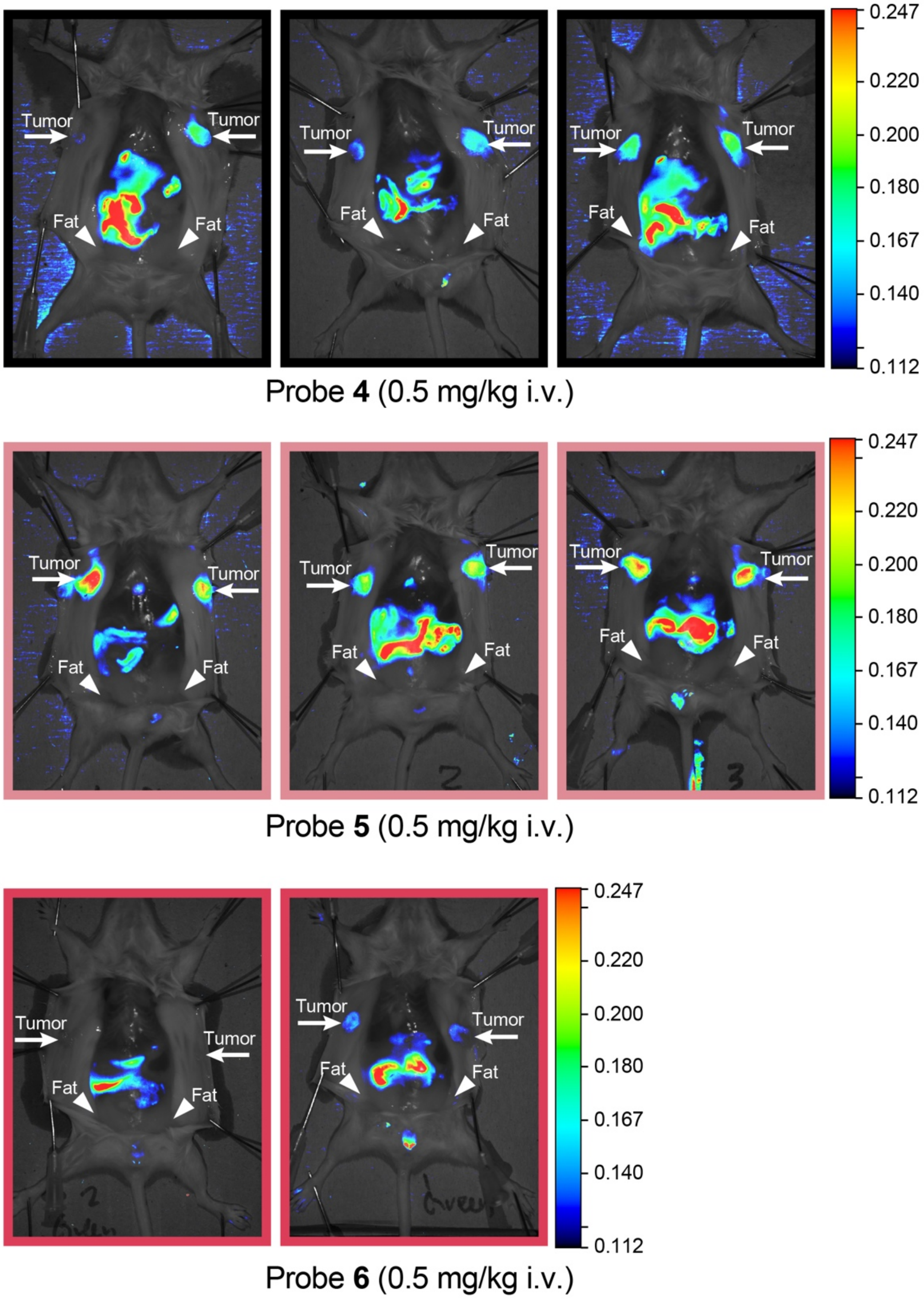


**Fig. S7: Full set of images from imaging of breast tumors with optimized fluorescently quenched substrate probes.**
**(A)** Fluorescent signal in tumor bearing mice. Images of the full set of mice injected with probes **4-6** (0.5 mg/kg in 10% DMSO + 30% PEG-400, 3 h). Mice have been splayed to show mammary tumors. Location of tumors, and healthy fat pads are indicated.

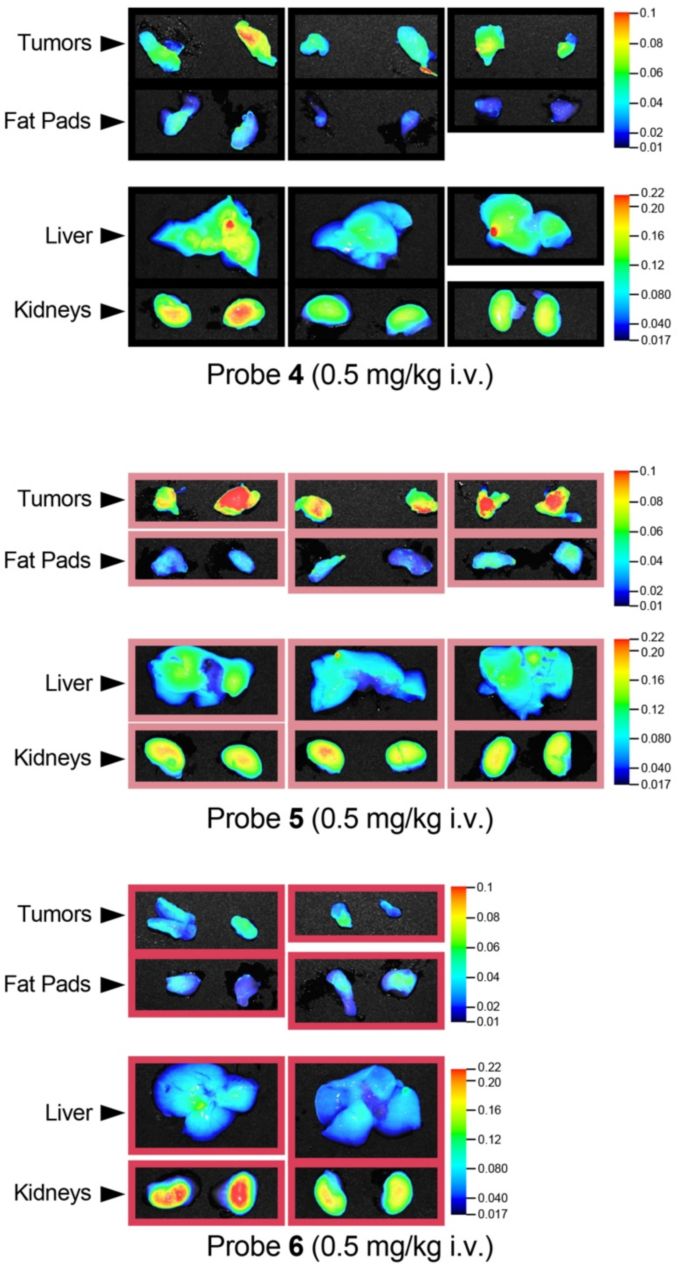


**Fig. S8: Full set of images for ex vivo analysis of fluorescence in organs in mice injected with optimized fluorescently quenched substrate probes.**
Fluorescent signal in tumor, fat pad, liver, and kidneys for animals injected with probes **4**-**6**.

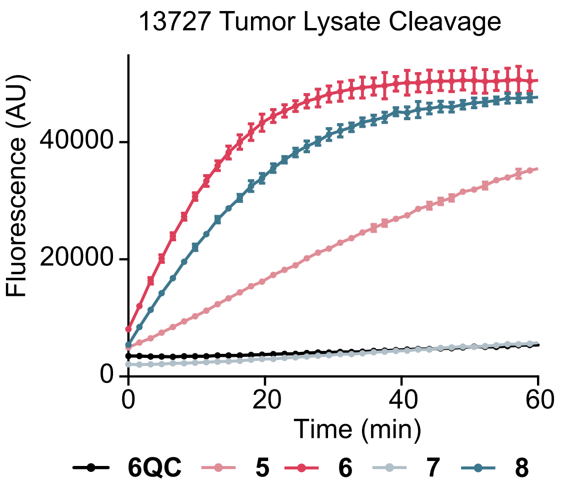


**Fig. S9: Cleavage of optimized probes in human breast cancer sample.** Progression curves for probes **4-8** in lysate of one representative human breast cancer sample. Plotted as Cy5 fluorescence over time. Values are plotted as mean value ± SD. n=3.
